# Evaluation of Engineering Potential in Undomesticated Microbes with VECTOR

**DOI:** 10.1101/2025.03.28.645341

**Authors:** Riley Williamson, Nicholas Dusek, Eglantina Lopez-Echartea, Megan K. Townsend Ramsett, Barney Geddes

## Abstract

Genetic engineering research has predominantly focused on well-characterized organisms like *Escherichia coli* and *Bacillus subtilis*, with methods that often fail to translate to other microorganisms. This limitation presents a significant challenge, particularly given the increasing isolation of large microbial collections through high-throughput culturomics. In response, we developed a scalable, high-throughput pipeline to evaluate the engineerability of diverse microbial community members we named VECTOR (**V**ersatile **E**ngineering and **C**haracterization of **T**ransferable **O**rigins and **R**esistance). We utilized a library of vectors with the Bacterial Expression Vector Archive (BEVA) architecture that included combinations of three antibiotic resistance genes, and three broad host range origins of replication (pBBR1, RK2 and RSF1010) or the restricted host range R6K with an integrative mariner transposon. We tagged each vector with green fluorescent protein and a unique nucleotide barcode. The resulting plasmids were delivered *en masse* to libraries of undomesticated microbes from plant microbiomes in workflows designed to evaluate their ability to be engineered. Utilizing OD_600_ and relative fluorescence measurements, we were able to monitor genetic cargo transfer in real time, indicating successfully engineered strains. Next-generation sequencing of plasmid molecular barcodes allowed us to identify specific vector architectures that worked well in particular bacterial strains from a large community. Modifications to the procedure facilitated isolation of engineered microbes. Our results underscore the potential of this approach to rapidly develop toolkits for the efficient engineering of a wide range of cultivatable microorganisms.

**Importance:** Undomesticated, cultured microbial strains contain a largely untapped reservoir of genetic potential for synthetic biology, and are increasingly being utilized in synthetic communities for microbial ecology research or biotechnology.. However, these strains often have unique physiological or ecological characteristics that make them difficult to engineer using traditional methods. Current approaches are often restricted by inefficient plasmid delivery and integration, which stifles progress in unlocking the promise of undomesticated strains. Our research addresses this challenge by developing VECTOR (**V**ersatile **E**ngineering and **C**haracterization of **T**ransferable **O**rigins and **R**esistance), a scalable, high-throughput pipeline that utilizes modular vectors and efficient engineering workflows to identify host range and improve plasmid uptake. By optimizing plasmid architectures and pooling them for simultaneous screening across a range of bacterial strains, VECTOR enhances engineering efficiency and opens new avenues for advancements in biotechnology.

## Introduction

The challenges and opportunities of modifying undomesticated microbial strains have become a focal point in genetic engineering. Organisms like *Escherichia coli* and *Bacillus subtilis* can be unsuitable for synthetic biology applications depending on the desired product (Goeddel *et al*., 1979; Hwang *et al*., 2016) and exploring non-model bacterial chassis is crucial to overcoming these limitations. Recent advances in culturomics and high-throughput techniques have opened the door to accessing a broader range of previously uncultivated microbes (Sarhan *et al*., 2019; Elmore *et al*., 2023). In many cases, wild microbes possess unique metabolic capabilities (Forum, 2013; Ata and Mattanovich, 2024), environmental resilience (Hirsch and Spokes, 1994; Barra, Danino and Garrido, 2020), and biochemical pathways that are highly desirable for biotechnological applications (Balagurunathan *et al*., 2022). Furthermore, synthetic communities constructed of cultivated microbes have emerged in microbial ecology as powerful simplified approaches to investigate causality in microbiome research (Vorholt *et al*., 2017). Genetic engineering in synthetic communities can facilitate advanced strain tracking methods (Daniel *et al*., 2024; Jorrin *et al*., 2024; Ordon *et al*., 2024), or may be required to test hypotheses about gene function derived from their study.

Undomesticated strains often lack the genetic tools necessary for advanced engineering. Their genetic stubbornness and metabolic complexity make traditional approaches slow and labor-intensive (Brophy *et al*., 2018). Overcoming these barriers has been further hindered by the absence of advanced, high-throughput genome engineering tools tailored for non-model microbes (Gilbert *et al*., 2023a). The genetic backgrounds and intracellular conditions of these strains often demand bespoke solutions to fully exploit their potential (Elmore *et al*., 2023). Modular vectors have been designed for efficient and scalable genetic modification across a range of microbial species (Silva-Rocha *et al*., 2013; Geddes, Mendoza-Suárez and Poole, 2019; Martínez-García *et al*., 2023; Geddes *et al*., 2024).

A majority of current plasmid-based engineering methods, while promising in their controlled delivery of genetic cargo, face limitations that restrict their broader application. A plasmid is not a “skeleton key” that fits all microorganisms; its design often needs to be tailored to specific hosts. This underscores the importance of scalable techniques to identify key plasmid components for successful engineering across diverse species. High-throughput approaches to explore engineering space, such as those developed for *Corynebacterium glutamicum* or *Psuedomonas putida* (Calero, Jensen and Nielsen, 2016; Tenhaef *et al*., 2021), have improved efficiency but focus on single strains. Recent innovations, like the MAGIC system by Ronda et al. or the POSSOM system by Gilbert et al., are pioneering approaches for community engineering and demonstrate the potential of using modular plasmids, fluorescent markers, and nucleotide barcodes to track and optimize engineering efforts across complex microbial communities (Ronda *et al*., 2019; Gilbert *et al*., 2023, 2024).

By advancing these high-throughput methods and tools, our research aims to unlock the genetic potential of undomesticated microbial strains and pave the way for more efficient and scalable engineering across communities of cultivated microorganisms. This study aims to: (1) investigate the success of plasmid-based engineering in diverse microbes via optical density, fluorescence, and next generation sequencing (NGS), and (2) facilitate the efficient isolation of engineered microbes. To do this we developed VECTOR **(V**ersatile **E**ngineering and **C**haracterization of **T**ransferable **O**rigins of Replication and Antibiotic **R**esistance), a new approach for microbial engineering focused on throughput and scalability by incorporating an acoustic liquid handling robot. For ground-truthing of our approaches, we applied them to strain libraries from plant microbiomes.

## Methods

### Strain Growth and Selection

A variety of bacterial strains were used for cloning purposes in this study (Supplemental Table S1). *E. coli* strains were grown at 37°C in liquid or solid Luria-Bertani (LB) medium and supplemented with 10μg/ml delta-aminolaevulinic acid (ALA) when working with *E. coli* ST18. When selecting for transformed cells containing the *lacZ* operon, X-gal was added at a concentration of 40μg/ml. Antibiotics were added to media, when necessary, at the following concentrations: gentamicin 10μg/ml, kanamycin 25μg/ml, neomycin 200μg/ml, spectinomycin 50μg/ml, ampicillin 100μg/ml, and tetracycline 5μg/ml. Our mock conjugation trials used well characterized nitrogen-fixing bacteria (Supplemental Table S1), grown in TY media with 0.5M CaCl_ at 28°C. For the corn and pea engineering trials, 94 culturable isolates with unique V4 16S rRNA gene sequences (47 from corn and 47 from pea) were selected from a culturomics study (Supplemental Table S2) (Lopez-Echartea *et al*., 2025). Corn and pea microbiome isolates were grown at room temperature or 28°C. Various concentrations of antibiotics were used for selection of transconjugants, and these concentrations have been noted in the descriptions of experiments.

### Golden Gate Cloning of Plasmid Library

BsaI or BsmBI Golden Gate assembly of vectors and plasmids was performed using the NEBridge® Golden Gate Assembly Kit (BsaI-HF®v2) or (BsmBI-v2) as recommended by the manufacturer (New England Biolabs).

We incorporated mariner transposon elements into a BEVA vector by cloning the mariner C9 transposase, an ampicillin resistance gene, and R6K origin of replication together, with terminal mariner mosaic elements. Following the BEVA assembly, a Golden Gate cloning site and antibiotic resistance cassette were incorporated into the transposable element generating pNDGG071.Two regions from pMarC9-R6K were amplified using primers RWND005/006 and RWND007/008 (Supplemental Table S3). Primers were designed to modify an endogenous BsmBI site through a single nucleotide change and to incorporate new BsmBI sites for compatibility as a BEVA “Position 3” module (Geddes et al., 2019). Products were cloned into the pGGAselect backbone using BsaI Golden Gate assembly and transformed into *E. coli* DH5α. Successful cloning was validated by restriction digest. Next, BsmBI Golden Gate assembly was used to construct R6K-based mariner transposon plasmids using various BEVA modules (Supplemental Table S4). Three variants with gentamicin, neomycin, or tetracycline resistance were assembled by combining pNDGG071 with pOGG004 (Level 1 Golden Gate cloning site), and either pNDGG001, pOGG009, or pOGG042 (antibiotic resistance modules). Products were electroporated into *E. coli* EC100D™ Pir+ and validated by whole plasmid sequencing by Plasmidsaurus using Oxford Nanopore Technology with custom analysis and annotation.

Our plasmid library was constructed via BsaI Golden Gate cloning, utilizing the three mariner vectors described above and nine BEVA2.0 vectors obtained containing the origins RK2, pBBR1, and RSF1010 (Geddes *et al*., 2024). Vectors were combined with an open reading frame expressing green fluorescent folding protein (sfGFP) and one of twelve oligo barcodes with Plasmid-ID architecture (Figure 1) (Supplemental Table S4 and S5) (Mendoza-Suárez *et al*., 2020). BEVA2.0 derivative plasmids were isolated by transforming into *E. coli* ST18, fluorescent colonies were selected, and plasmids were validated via restriction digest. Due to concatomerization observed during cloning directly into ST18, mariner-derived plasmids were alternatively verified first in *E. coli* EC100D™ Pir+ before being transfer to ST18.

**Figure 1.**
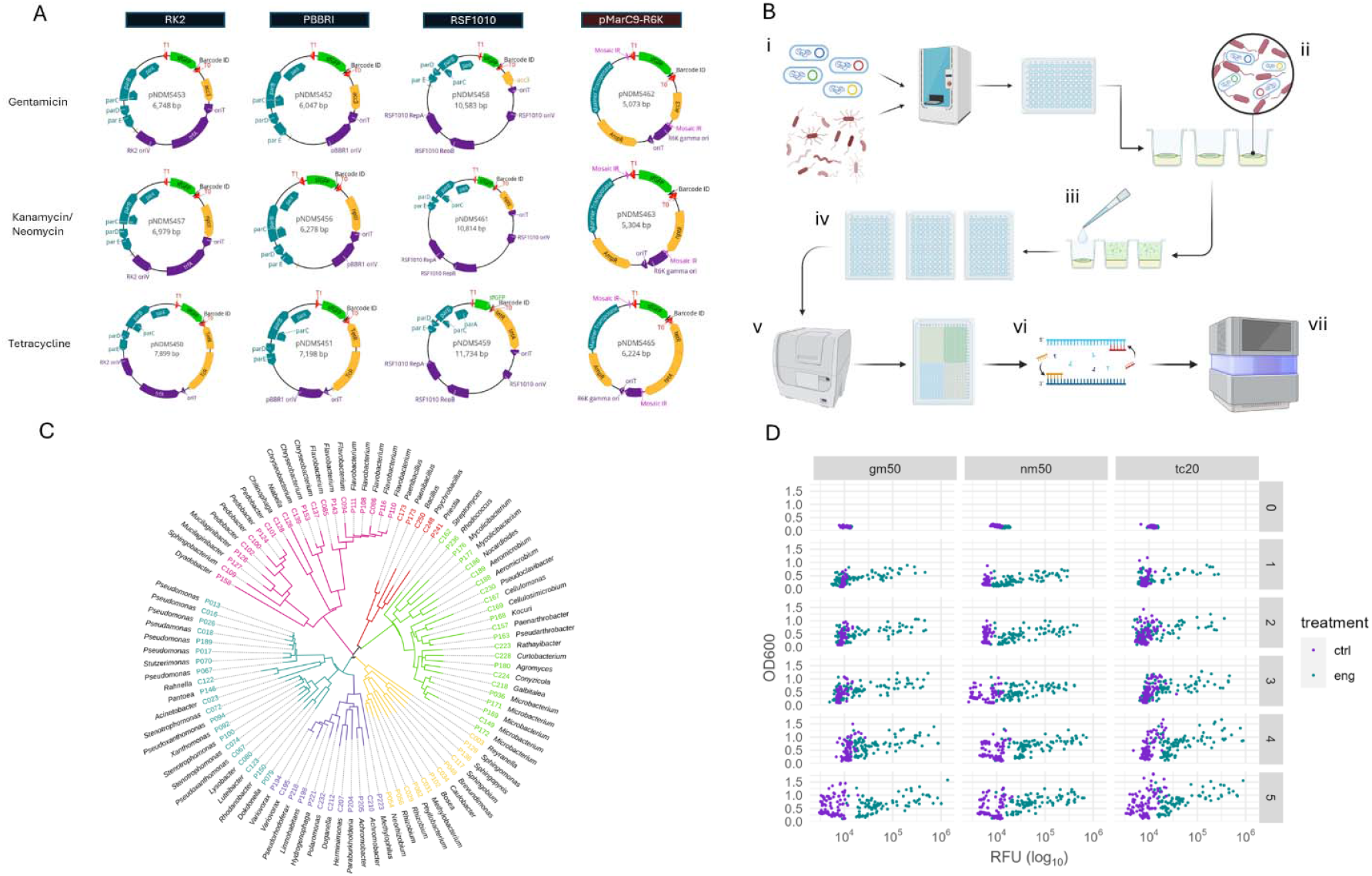
Design and testing of VECTOR^L^ workflows for screening plasmid-based engineering success in diverse microbes. **(A)** Plasmid suite for tracking microbe engineerability, constructed using BEVA architecture. Each column denotes the vector’s origin of replication and each row, its antibiotic resistance module. **(B)** VECTOR^L^ engineering workflow. (i) Donor cells containing barcode tagged conjugative plasmids are pooled together and transferred to mating spots along with recipient cells. (ii) Mating spots incubated. Each spot containing a single recipient strain and the entire plasmid library. (iii) Post-incubation, mating spots are resuspended. (iv) Resuspensions are added to various selection plates containing liquid media, supplemented with various antibiotics. Media also lacks additives to select against auxotrophic donor cells. (v) Selection plates are measured daily, using a plate reader to monitor growth and fluorescence. (vi) Selection plates undergo a 2-step barcoding PCR. PCR1 adds indexing primers to identify each well in a 96-well plate. Wells are then pooled together and PCR2 incorporates indexing primers to denote each plate. (vii) Amplified DNA is sequenced using Illumina technology. **(C)** Maximum likelihood phylogenetic tree displaying the relationship among the 94 corn and pea derived strains used in our study, based on the V4 16S. Strains are labeled with their ASV number and Genus. Strains are color coded by class: *Cytophagia* (pink), *Bacilli* (red), *Actinomycetia* (green), *Alpha-proteobacteria* (yellow), *Beta-proteobacteria* (purple), and *Gamma-proteobacteria* (turquoise). (D) Dot plot grid containing 18 individual plots, each displaying log_10_(RFU) on the x-axis and OD_600_ on the y-axis. The grid is organized with rows representing days 0–5 and columns corresponding to the antibiotics used for selection. Points are color-coded: engineered strains are shown in turquoise, control strains in purple.

### High Throughput Conjugations with Liquid Media Selection

The protocol for high throughput conjugation with liquid media selection (VECTOR^L^) has been deposited at DOI: dx.doi.org/10.17504/protocols.io.q26g7m581gwz/v1 and summarized below. TY broth (150μl/well) was inoculated with recipient cells in a 96-well plate, sealed, and incubated at room temperature for 3–7 days, with growth monitored regularly. To normalize growth across the library, recipient cultures at OD_600_ ∼0.4–2 were subcultured (15μl into 135μl TY) and incubated overnight at 28°C with orbital shaking set to 215rpm, while donor cultures were grown overnight in LB+ALA at 37°C, also shaking at 215rpm. OD_600_ of overnight cultures was measured with a BioTek Cytation 5. Donor and recipient cells were combined in 96-well mating spots using an Echo-525 acoustic liquid handling robot based on the Echo Plate Reformat programs “CornandPeaTrial_Cherrypick.xlsx” or “MockConjugation_Cherrypick.xlsx” (Supplemental Table S6). Cell volumes were calculated to add a total of ∼2 million cells/μl, within a 5μl mating spot, assuming an OD_600_ measure of 1.0 contained ∼100 million cells/μl. Conjugation plates were incubated overnight at 28°C, and mating spots were resuspended in 150μl TY broth. Resuspensions were added to selection plates at a 1:20 dilution in 150μl TY + antibiotics and stored in the dark at room temperature for five days. OD_600_ and RFU (485/20nm excitation, 528/20nm emission) were measured daily using the Biotek Cytation 5. On day 5, 10μl of each culture was transferred to a PCR plate for storage at -20°C for further NGS, and the remaining culture was combined with glycerol (40% final concentration) and stored at -80°C for longer term storage. OD_600_ and RFU data were compiled in Microsoft Excel and analyzed using custom scripts “Growth_Curve_Analysis.R”, “Growth_Curve_Analysis_Engineered.R”, “MockConjugation_OD_GrowthCurve.Rmd”, and “MockConjugation_RFU_GrowthCurve.Rmd”(Supplemental Table S6, https://github.com/NDSU-Geddes-Lab/eng-potential-vector).

### High Throughput Conjugations with Solid Media Selection

The protocol for high throughput conjugation with solid media selection (VECTOR^S^) has been deposited at http://dx.doi.org/10.17504/protocols.io.q26g7m581gwz/v1 and summarized below.

Conjugations selected on solid medium followed the same steps detailed above, using the “CornandPeaTrial_SinglePlasmid_Cherrypick.xlsx” or “MockConjugation_MultiPlasmid_Cherrypick.xlsx” procedures for mating spot preparation in the Echo-525 (Supplemental Table S6, https://github.com/NDSU-Geddes-Lab/eng-potential-vector). After conjugation, mating spots were resuspended in 150 µl TY broth and transferred to a sterile 96-well plate. Cultures were serially diluted in saline solution? from 10□^1^ to 10□ □.

Dilutions were plated onto TY agar + antibiotic as an array of 10 µL spots in single-well rectangular plates, dried under a hood, sealed, and stored in the dark at room temperature for five days. On day five, plates were imaged with a BioTek Cytation 5 using 1.25X PL APO magnification, with FDP fluorescence imaged at 469 Excitation, 525 Emission (LED 10, 5 ms integration time, 17.6 gain) in a 9×13 montage with auto tile overlap.

Streak purified engineered *S. meliloti* colonies were screened for their possession of pNDMS451 with long read whole-genome sequencing by Plasmidsaurus using Oxford Nanopore Technology.

### Barcode Sequencing of Plasmids Following Liquid Media Selection

For DNA extraction, the 10 µl of each conjugation culture,transferred to a 96-well PCR plate at the end of VECTOR^L^, was mixed with 14 µl alkaline lysis buffer (5 ml 20% SDS, 2 ml 10 mM NaOH, 93 ml water, pH 12), incubated at 95°C for 20 minutes, neutralized with 14 µl neutralization buffer (1.936 g of Tris-HCl, 400 mL of water, pH 7.5), and stored at -20°C. A dual indexing strategy was used for next-generation plasmid sequencing. Primary PCR added dual well-ID barcodes using a set of 12 forward and 8 reverse primers that amplify the Plasmid-ID (Supplemental Table S7), and secondary PCR added Nextera barcodes and Illumina adaptors (Supplemental Table S8) (Supplemental Figure S1A) (Schumacher *et al*., 2025). PCRs were prepared using KAPA Hifi HotStart ReadyMix (Roche) in miniaturized reactions with 0.5µl of template DNA and totaling 5µl per reaction. PCR samples were carried out in an Echo-525 Acoustic handling robot (Benz *et al*., 2024). The primary PCR cycling parameters were 95°C for 3 minutes, 25 cycles of 95°C for 30 seconds, 62°C for 30 seconds, 72°C for 30 seconds, and 72°C for 5 minutes. The secondary PCR was performed in with the same parameters for 8-12 cycles. PCR products were purified after each step with AMPure XP beads according to manufacturer’s recommendations. The final products were quantified with Qubit, and pooled in equimolar ratios for sequencing with an Illumina Miseq. Following sequencing, Nextera barcodes were demultiplexed by the sequencer to separate reads into bins corresponding to source plates (Supplemental Figure S1B). Reads were further demultiplexed to represent the portion of Plasmid IDs in each well using a custom script (Schumacher *et al*., 2025), https://github.com/NDSU-Geddes-Lab/plasmid-id). Reads were then assigned to each plasmid backbone according to the Plasmid-ID and then normalized based on anticipated plasmid copy number: 7 for RK2 (Kües and Stahl, 1989), 30 for pBBR1 (Jiang *et al*., 2017), 12 for RSF1010 (Meyer, 2009), and 1 for pMarC9-R6K transposon (**Supplementary Methods)**.

### Fluorescence and Plasmid Frequency Validation

As a part of the ground truthing process, glycerol stocks generated in the high throughput liquid conjugation protocol were streak-plated onto TY agar + antibiotic, and each parental strain on an antibiotic free plate, and stored in the dark at room temperature for up to five days. The presence of sfGFP fluorescence in each strain was visualized using a Bulldog Bio FastGene Blue/Green LED Transilluminator DE.

Single colonies from four specific strains (C162 (Tc), P189(Tc), C169 (Nm), and P150 (Tc)) were used for verification of plasmid contents by colony PCR. PCR was performed using primers specifically designed to bind regions unique to each plasmid *ori* variant, enabling identification of the *ori* present in each colony (Supplemental Table S3). Each 22.5 μl PCR reaction contained 2.5 μl DNA, 2.5 μl OneTAQ Buffer (New England Biolabs), 0.5 μl dNTPs 0.2mM (ThermoFisher), 0.5 μl of each primer 1 µM, 0.125 μl TAQ polymerase (New England Biolabs and 15.875 μl water. The cycling conditions included an initial denaturation at 94°C for 2 minutes, followed by 32 cycles of 94°C for 30 seconds, 54°C for 30 seconds, 72°C for 30 seconds, and a final extension at 72°C for 5 minutes. PCR products were then run on an 1.5% electrophoresis gel to verify band size.

### Logistic regression modeling of engineerability

To support future high-throughput screening of transconjugants for verification of engineerability, we fit several logistic regression models using the glm function from the stats package in R (R Core Team, 2024), with OD, RFU, and taxonomic class as predictors and the ground truth labels of engineered vs. non-engineered (within each antibiotic condition) as the response. Taxonomic class of each strain was treated as a single factor with 6 levels (Figure 1C). OD_600_ and RFU, measured daily from 0 to 5 days, were transformed to ΔOD and ΔRFU as defined below, where OD_*t*_/RFU_*t*_ represents the measurement at day *t*.

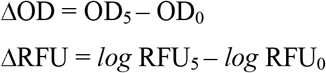

In total, 6 logistic regression models were developed: 2 models per antibiotic (OD/RFU and taxonomic class) for each of the three antibiotic treatments (gentamicin 50μg/ml (Gm50), neomycin 50μg/ml (Nm50), and tetracycline 20μg/ml (Tc20)), according to the general model formulae below.

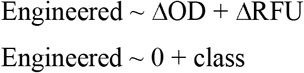

For the models using taxonomic class as the predictor, the intercept was set to 0 so that an explicit effect would be included for each class (as opposed to the default behavior of glm, where the first class is set as the reference level). Each model was assessed for goodness-of-fit by a χ^2^ test on the residual deviance and by visual assessment of the residuals (**Supplemental Methods**).

## Results

### Plasmids to facilitate high-throughput engineering evaluations

An overall objective in this research was to develop a high-throughput method for screening the engineering capabilities of large microbial communities *en masse*. To facilitate such a method, we utilized a library of plasmids that we expected to function in a wide variety of microbes and endowed measurements of engineering success via sfGFP fluorescence and NGS of DNA barcodes. As a platform, we used a suite of recently developed BEVA2.0 plasmids that included several broad host-range origins of replication (pBBR1, RSF1010 and RK2) and antibiotic resistance cassettes (gentamycin, neomycin and tetracycline) in each combination, with a MoClo BsaI Golden Gate cloning site (Weber *et al*., 2011; Geddes, Mendoza-Suárez and Poole, 2019; Geddes *et al*., 2024). We further expanded the library by developing a set of plasmids for transposition (pMarC9-R6K) that included a backbone with an R6K origin of replication, ampicillin resistance, and a mariner C9 transposase and a transposable element flanked by mariner mosaic ends. These conserved the BEVA architecture by incorporating the same MoClo BsaI Golden Gate cloning site and antibiotic resistance modules as the BHR plasmids into the transposable element. Each of the twelve resulting plasmids was used for the introduction of sfGFP preceded by J23106 (promoter) and B0032m (ribosome binding site) expression elements from CIDAR (Iverson *et al*., 2016), and followed by a DNA molecular barcode with Plasmid-ID architecture (Figure 1A, Supplemental Table S5) (Mendoza-Suárez *et al*., 2021).

### A screening pipeline to evaluate engineering success in diverse microbes

We devised VECTOR, an efficient workflow to introduce the above plasmid suite *en masse* into diverse recipient strains by conjugation, utilizing growth (OD_600_), sfGFP fluorescence, and Plasmid-ID NGS to evaluate engineering success following antibiotic selection (Figure 1B). The workflow was performed as follows:

#### Step 1: Donor and recipient cells are combined in a miniaturized mating spot by acoustic liquid handling robot

We used ST18 as the donor to facilitate counterselection based on L-auxotrophy. A library of donors, each containing one of the 12 plasmids above, was pooled in equal proportion. The pooled donor cells were combined with an array of recipient strains; the targets for assessment of engineering potential. Donors and recipients were also added individually to control wells, with each treatment combination above performed three times independently.

#### Step 2: Liquid selection of transconjugants

Following incubation of the mating spot, spots were resuspended in liquid media containing tetracycline, neomycin or gentamicin and deposited into 96-well plates. Selection occurred during 5 days of growth in the presence of antibiotics.

#### Step 3: Monitoring engineering success by growth and fluorescence

Each day throughout the selection outgrowth, 96-well plates were imaged using a plate reader and data for growth (OD_600_) and GFP fluorescence (RFU) were collected.

#### Step 4: Assessment of individual plasmid success by NGS

Following selection, an aliquot from each well was used for library preparation and NGS. We determined the relative abundance of Plasmid-ID barcodes in each well, and used this data to assess biases in successful engineering by different plasmid architectures from the initial pool.

### Workflow optimization with a domesticated strain set

We optimized the VECTOR^L^ protocol by applying it to a set of six commonly studied laboratory strains; diazotrophs from the plant microbiome (*Sinorhizobium meliloti* RmP110, *Mesorhizobium loti* R7A, *Azospirillum brasilense* Sp7, *Rhizobium spp*. IRBG74, *Klebsiella oxytoca* M5A1, and *Azorhizobium caulinodans* ORS571). Pilot experiments evaluated recipient-to-donor ratios (1:1, 2:1, 5:1) and three different concentrations of each antibiotics (Gm 20 μg/mL, 50 μg/mL and 100 μg/mL; Nm 20 μg/mL, 50 μg/mL and 100 μg/mL; Tc 5 μg/mL, 10 μg/mL and 20 μg/mL). We evaluated engineering success of the target strains based on the difference between growth and fluorescence in engineered wells compared to control wells, and found that changing the recipient-to-donor ratios had limited effect on outcomes, though the 5:1 ratio was more successful in some cases (Supplemental Figure S2A). We identified antibiotic concentrations which maximized the difference in growth between control and engineered wells (Supplemental Table S9). Subsequent implementations of the protocol were performed at these antibiotic concentrations (Gm 50 μg/mL, Nm 50 μg/mL and Tc 20 μg/mL), and a recipient-to-donor ratio of 5:1. NGS of Plasmid-IDs revealed a notable presence of each plasmid-type (pBBR1, RK2, RSF1010 and pMar9-R6k), with their relative read abundance correlating to expected copy number and limited differences between strains (Supplemental Figure S2B).

### Evaluating engineering potential in environmental isolates from plant microbiomes

Next, we investigated the utility of VECTOR^L^ for exploring the engineering potential in collections of environmental isolates. We used 94 members from a pea and corn root microbiome collection developed by high-throughput culturomics (Lopez-Echartea *et al*., 2025). These members represented undomesticated isolates from the plant microbiomes belonging to *Cytophagia, Bacilli, Actinomycetia, Alpha-proteobacteria, Beta-proteobacteria*, and *Gamma-proteobacteria* (Figure 1C). The 12-plasmid library was introduced to each member by miniaturized conjugation and the selection was performed concurrently in liquid medium containing Gm 50μg/mL, Nm 50μg/mL or Tc 20μg/mL. When growth and fluorescence were measured across five days of selection in each antibiotic, we observed a distinct clustering of these values in transconjugant strains relative to the control wells that continued to increase across the five days of selection period (Figure 1D, Supplemental Figure S3). Fluorescence measurements were the most effective at separating the treatments, with growth measurements presumably obscured by high growth of control wells in strains displaying resistance to antibiotic-selection. NGS data of the barcodes from this dataset also allowed us to visualize the distribution of plasmid reads in each strain, and explore any biases in plasmid success (Supplemental Figure S4 and S5). We found several examples wherein reads for a specific plasmid or plasmid-type (based origin of replication) significantly exceeded the others (Fig or table).

### Ground-truthing predictions of engineering success

To explore the reliability of growth/fluorescence measurements to predict engineering success, we streak-plated one representative of each strain per antibiotic combination, and evaluated its engineer success based on the isolation of single colonies that matched colony morphology to the expected recipient strain and showed sfGFP fluorescence (Figure 2A). Overall, we found evidence for successful engineering of 55/94 strains with at least one antibiotic, with engineered representatives widely distributed across the total phylogeny of the strains. Of the engineerable strains, about half (28/55) were successfully engineered with all three antibiotics, with the remaining failing to be engineered with one or two antibiotics. We used these data to develop logistic regression models that evaluated the predictability of engineering success in each antibiotic based on the difference in OD_600_ or RFU measuremenents between experimental and control wells (Figure 1D) and taxonomic class (Figure 2B). All three models using ΔOD_600_ and ΔRFU as predictors were considered good fits to the data by the χ^2^ test (*p* > 0.10) (Table 1, Figure 2C-E), whereas the three models using taxonomic class were not good fits (Supplemental Data). Comparing the OD_600_ and RFU data, *p*-values for the effects of ΔOD_600_ and ΔRFU were statistically significant (*p* < 0.05) across all three antibiotics. For successful models, group separation (engineered vs. non-engineered) is further illustrated in Figure 2C-E, which largely corroborates the overall model fits and effect sizes for the three different antibiotics, with Gm50 showing the greatest separation, followed by Nm50, and lastly Tc20.

**Table 1.** Model coefficients and *p*-values for logistic regression models on ΔOD and ΔRFU. Overall model *p*-values are from the χ^2^ goodness-of-fit test (upper tail). Coefficients are reported as the original *log* odds from glm, with the standard odds (exponentiated *log* odds) and *p*-value in parentheses. In logistic regression, coefficients represent the *log* odds of a positive result over a negative result for each unit increase in a continuous predictor. When assessing model fit on the whole, failure to reject the null for the χ^2^ test (typically at *p* > 0.05) indicates that the residual deviance follows the expected distribution, meaning the model can be considered a good fit to the data. Labels at the top of each column refer to the antibiotic and concentration used: gentamicin 50μg/mL (Gm50), neomycin 50μg/mL (Nm50), and tetracycline 20μg/mL (Tc20).

| | **Gm50**<br>($p = 0.908$) | **Nm50**<br>($p = 0.509$) | **Tc20**<br>($p = 0.390$) |
| --- | --- | --- | --- |
| **Intercept** | -3.3090<br>(0.0366, $p = 0.000155$) | -1.9988<br>(0.136, $p = 0.000736$) | -2.9753<br>(0.051, $p = 0.000006$) |
| **$\Delta OD$** | 4.6084<br>(100.322, $p = 0.000119$) | 3.0850<br>(21.867, $p = 0.000482$) | 1.7872<br>(5.973, $p = 0.0322$) |
| **$\Delta RFU$** | 2.8642<br>(17.536, $p = 0.000418$) | 1.7981<br>(6.038, $p = 0.010100$) | 3.2077<br>(24.721, $p = 0.000038$) |

**Figure 2.**
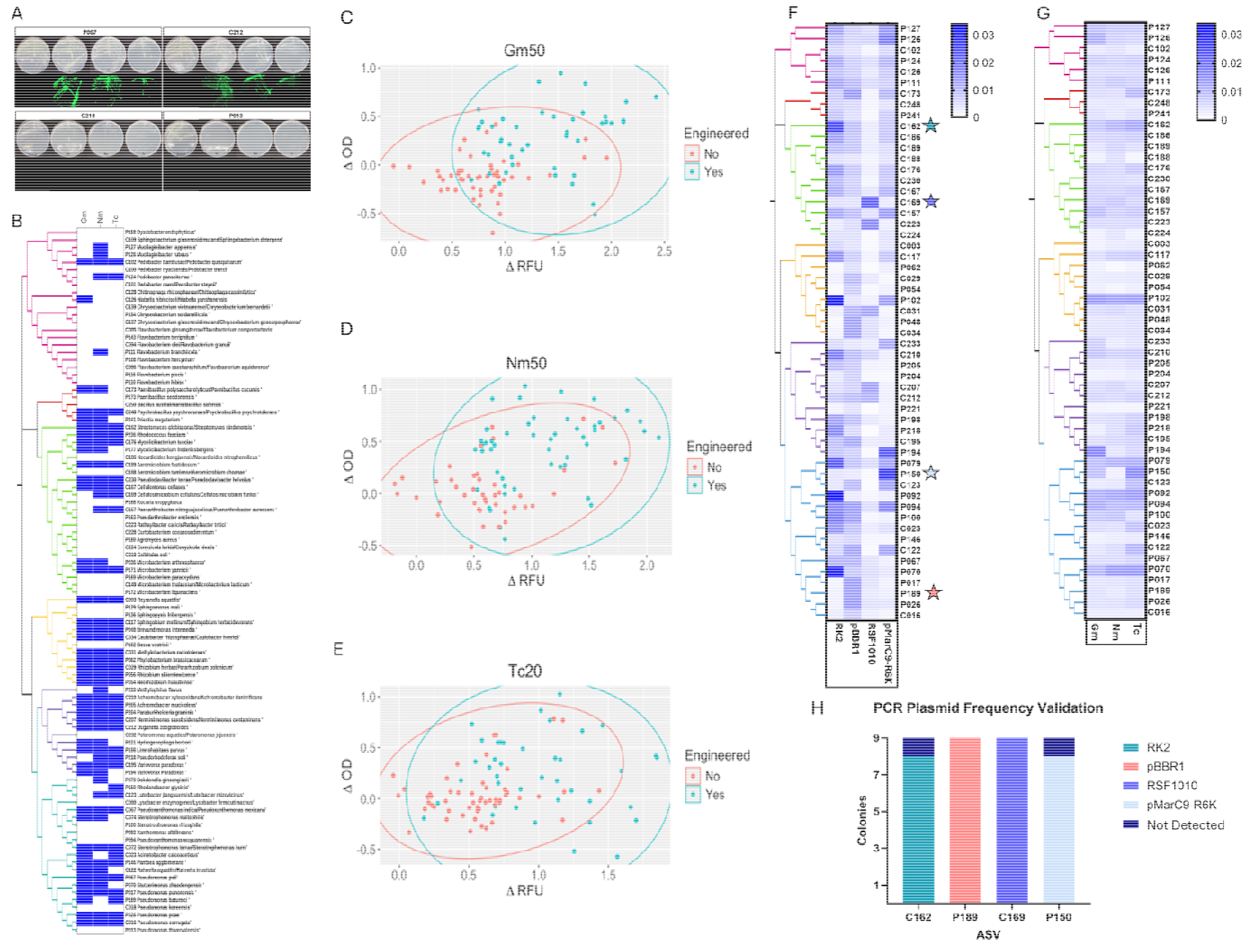
Verification of VECTOR engineering events. **(A**) Representative streak plates demonstrating strain growth on selective antibiotics, with comparisons to the parental strain based on colony morphology (top) and GFP fluorescence (bottom). **(B)** A conditions table showing the engineering success of each strain (listed on the right) under specific antibiotic conditions (labeled at the top). A strain was classified as engineered if it exhibited visible colony growth matching the parental strain and expressed fluorescence. Phylogenetic relationships, based on maximum likelood of the strains’ V4 16S,are displayed on the left side of the table. **(C-E)** Dot plots depicting cell growth and fluorescence at the end of day 5. Each plot displays ΔRFU (log10) on the x-axis and ΔOD_600_ on the y-axis. Plots are organized by antibiotic condition: (C) gentamicin (50 μg/ml), (D) neomycin (50 μg/ml), and (E) tetracycline (20 μg/ml). Points are color-coded by their engineering success observed in streak plating: blue for engineered strains and red for non-engineered strains. Ellipses are plotted around each group (engineered and non-engineered) to better indicate group separation. ΔRFU and ΔOD_600_ values were calculated by subtracting control values for each strain. **(F&G)** Heat maps displaying the relative read counts of all plasmids across the tested bacterial strains, with the darker blue representing a high number of reads. Reads have been normalized to account for the copy number of each plasmid. Squares on the heat maps are plotted according to relative abundance amongs all reads. Heat map (F) organizes the data by plasmid *oris*, while heatmap (G) organizes the data by antibiotic conditions, listed at the bottom of each column. The phylogenetic relationships of the strains are shown on the left side of each heatmap, and strain ASVs are noted to the right of each row. The intensity of blue shading within each square corresponds to plasmid read counts, with darker shades indicating greater plasmid abundance. Strains selected for PCR validation are marked with a blue star to highlight their inclusion in further analysis. **(H)** Plasmid frequencies based on presence observed in 9 colonies selected from four engineered strains. Plasmid origins are indicated by color.

Next, to ground-truth the reliability of NGS data of barcodes at predicting biases in plasmid engineering success, we used PCR screening to test the distribution of plasmid-types in one representative that we expected to primarily be engineered with each plasmid (C162 for RK2, P189 for pBBR1, C169 for RSF1010 and P150 for pMark-R6K) (Figure 2F). In each example, we found the expected plasmid to be the primary occupant of streak-purified colonies (Figure 2H).

Taken together, these data indicate this workflow can be used to predict 1) the ability to engineer microbes within a collection, and 2) specific plasmid architectures likely to be successful.

### Adjusted workflows for purification of targeted engineering events

To facilitate the isolation of desired engineering events we also developed an adjusted protocol, VECTOR^S^, that focused on isolation of engineered microbes as single colonies following solid-media selection (Figure 3A). We tested two schemes: (1) employing multiple plasmid variants to engineer a single strain. (2) using a single plasmid variant to engineer multiple bacterial strains. Modifications to the workflow presented above included solid media selection using drop-plates in serial dilution, and screening of engineering events by imaging solid agar plates and identifying fluorescent single colonies.

**Figure 3.**
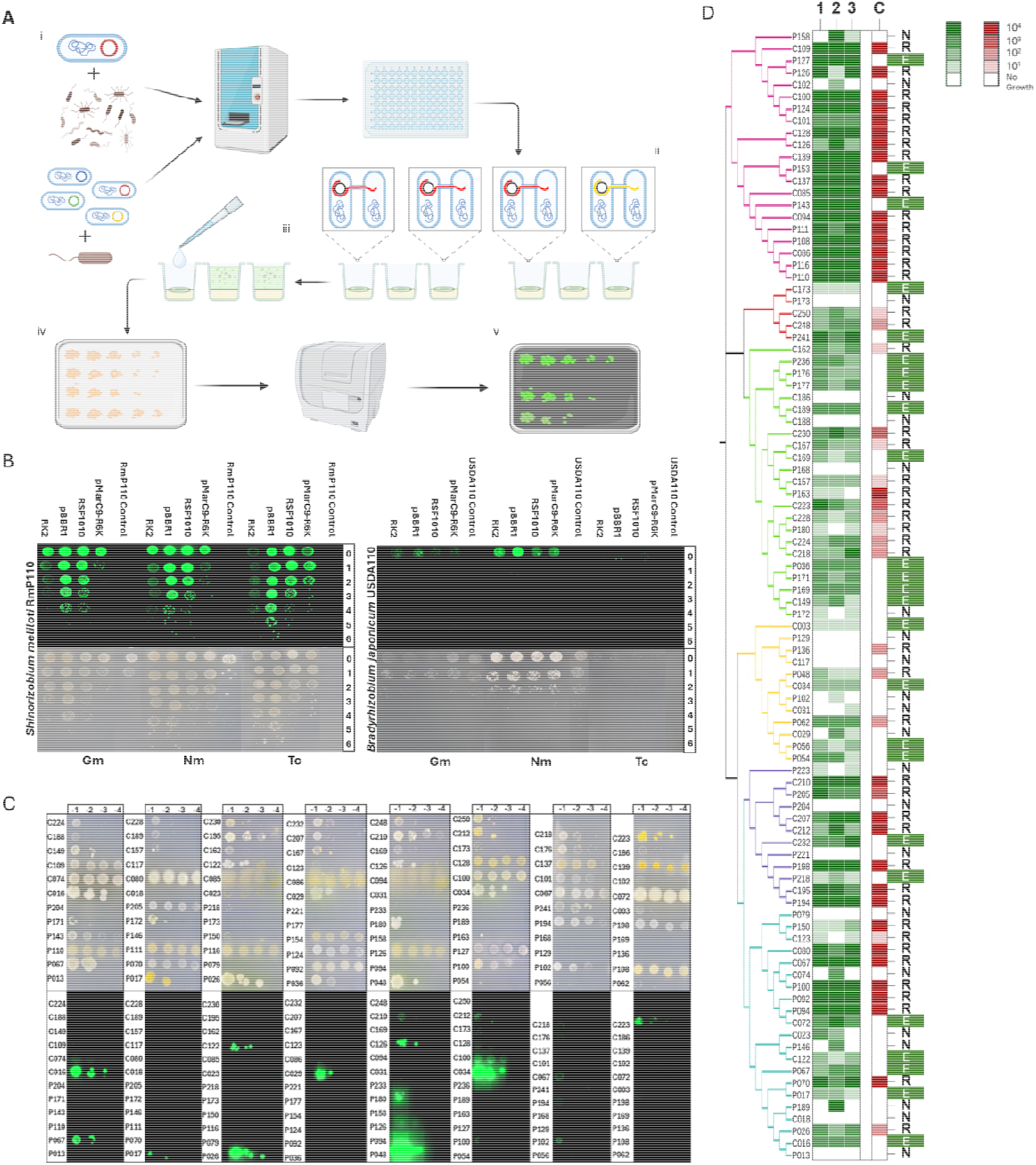
Targeted engineering with VECTOR^S^. **(A)** VECTOR^S^ engineering workflow (i) Donor cells containing the same barcode tagged conjugative plasmid variant are transferred individually to mating spots along with various recipient cells (top). Donor cells containing various barcode tagged conjugative plasmids are transferred individually to mating spots along with recipient cells of the same strain (bottom) using an acoustic liquid handler. (ii) Mating spots are incubated. Each spot containing one of the recipient strains and a single member from the plasmid library (left) or a single recipient strain and one of many members from the plasmid library (right). (iii) After incubation, the mating spots are resuspended and subjected to serial dilution in saline solution. (iv) All dilutions are added to various selection plates containing solid agar media, supplemented with various antibiotics. Media also lacks additives to select against auxotrophic donor cells. Mating spot dilutions are added to selection plates containing the antibiotic that corresponds to the resistance module within the transferred plasmid. (v) Selection plates are monitored for several days as the spot dilutions grow. When proper growth is observed, the plates are imaged using a Cytation5 to identify colonies expressing GFP. **(B)** Spot plating results from multi-plasmid/single recipient engineering trials. Images of plates for RmP110 and USDA110, showing both GFP fluorescence (top) and standard views (bottom) for the same plates. Plasmids are listed on the top of each image, organized by origin of replication. Antibiotic conditions and dilution levels are listed on the bottom and side of each image respectively. **(C**) Spot plating results from single plasmid/multi-recipient engineering trials, continued. The image contains one of the triplicate plates for the corn and pea community trial, showing both standard views (above) and GFP fluorescence (below) for the same plates. The image has been split to label each strain, to the left of each spot, and the dilution level, listed above. **(D)** Spot plating results from single plasmid/multi-recipient engineering trials. Heat maps representing triplicate studies for each corn and pea ASV, with ASVs listed to the left of each row. These heat maps show the dilution levels at which consistent growth was observed for each plasmid, green for the engineered cells and red for their controls. Replicate numbers and controls are indicated at the top of each heat map. The final column on the right of each map displays the engineerability designation: E for Engineered, N for Not Engineered, and R for Resistant.

We evaluated the ability of each plasmid in our library to engineer a readily engineerable strain (*S. meliloti* RmP110) and a widely considered challenging-to-engineer strain (*Bradyrhizobium japonicum* USDA110) using this approach (Medici *et al*., 2024). For each plasmid, on each antibiotic treatment in *S. meliloti*, fluorescent colonies were identified at high frequencies and able to be picked for streak purification, even despite growth of the control on the neomycin treatment (Figure 3B). As expected, engineering was much less efficient in *B. japonicum* which grew robustly on both neomycin and gentamicin treatments. Fluorescence was observed for pBBR1 in tetracycline, all four backbones in neomycin and for pBBR1 and RK2 in gentamicin. However, only tetracycline produced well isolated colonies due to background growth in the other antibiotics. We further streak purified engineered colonies in from the pBBR1 Tc treatment and verified the presence of the plasmid by long read whole-genome sequencing.

Next, we explored engineering many strains with a single plasmid by exposing the plant microbiome collection to a neomycin-based mariner plasmid (pNDMS463), relevant to the amenability of strains to high throughput mutant fitness profiling using RB-TnSeq (Wetmore *et al*., 2015). We followed the same approach, and despite a widespread variance in growth of the microbiome members on plates, we readily identified engineering events in isolated colonies based on fluorescence imaging (Figure 3C). We found successful engineering by the neomycin-based mariner plasmid in approximately ∼27% of the collection (25/94 strains). Growth of resistant controls was commonly observed in this setup (47/94 strains; ∼50%) and may have limited the isolation of engineering events.

Overall, adapting the high-throughput conjugation approaches to solid media and solid plate fluorescence screening enabled the recovery of engineered colonies from wild-type strains and diverse environmental isolates.

## Discussion

Here we developed and tested VECTOR, a set of workflows for evaluating plasmid-based engineering of diverse microbes by bacterial conjugation, and facilitating the rapid isolation of engineered strains from collections of undomesticated bacteria. We demonstrated utility of these workflows for engineering diverse microbiome isolates collected from crop plant roots by culturomics (Lopez-Echartea *et al*., 2025). In our tests, we successfully engineered at least 55 of 94 isolates. This included isolates distributed throughout the 6 classes including 16 orders tested, and we found that strain taxonomy was a poor predictor of engineerability (Supplemental Dataset B), though members of the classes *Cytophagia, Bacilli, Actinomycetia* were less readily engineered than the *Alpha-, Beta-* or *Gamma-proteobacteria* (**Supplemental Figure S4**).

The liquid selection-based VECTOR^L^ workflow facilitated monitoring of engineering in real-time through growth (OD_600_) and fluorescence measurements of vector-borne sfGFP (Figure 1D, Figure 2C-E). Logistic regression models of engineerability as a function of ΔOD and ΔRFU fit our data well, and all effects were statistically significant across the three different screened antibiotics. These results provide evidence that change in OD and RFU, with respect to the corresponding non-engineered control, have predictive value in the context of high-throughput screening (Table 1). Of course, the likelihood that a particular strain has been successfully engineered, on the basis of ΔOD and ΔRFU, depends on the magnitude of the difference between the engineered and non-engineered samples for each metric. For this reason, logistic regression models are particularly useful in that they can be applied to unknown samples to calculate a predicted likelihood (between 0 and 1) that a sample has been successfully engineered.

The use of a highly simplified sigma-70 promoter lacking regulatory elements from the Anderson promoter series from the Registry of Standard Biological Parts (https://parts.igem.org/Promoters/Catalog/Anderson) was presumably an important element of the widespread success of fluorescence engineering across the diverse bacterial taxa in our study. While promoters from this library have been tested robustly in *E. coli* and a few other organisms (Iverson *et al*., 2016; Grant, 2019; Liow, Go and Yew, 2019; Armetta *et al*., 2021), our data indicate they are widely functional in the bacterial kingdom, at least in the dominant taxa of plant microbiomes. In addition to facilitate ground-truthing of successful engineering events by VECTOR^L^ (Figure 2AB), the fluorescence module allowed us to distinguish engineered colonies from background growth in the solid-media VECTOR^S^ for isolation and purification (Figure 3BC).

The incorporation of Plasmid-ID sequencing into VECTOR^L^ allowed us to measure the relative success of different plasmid architectures in individual isolates (Figure 1E, Figure 2 F-G). We found these predictions to be reliable at identifying strains that were predominantly engineered by one plasmid-type (Figure 2H). While all four plasmids showed some success across the phylogeny of tested strains, some trends for individual plasmid architectures performing better with particular lineages of microbes were apparent. For example, a bias towards RK2 success was apparent in members of the *Xanthomonadales*, whereas pBBR1 success was more common in *Pseudomonadales*. Still, there was widespread variation for individual success of plasmids among the isolates in different lineages; different members of *Rhizobiales, Burkholderiales* or *Micrococcales* were preferentially engineered with each of the four different plasmid origin types (Supplemental Figure 5).

The utility of concurrently screening diverse plasmids using VECTOR is highlighted by comparing the overall engineering success of VECTOR^L^ using the pooled plasmid library (55/94 strains, Figure 2B) to using VECTOR^S^ with only a single plasmid architecture (25/94 strains, Figure 3D). Incorporating more plasmid architectures could further increase engineering success, and we designed VECTOR to be highly scalable. Plasmid libraries can readily be expanded from our test collection of twelve to hundreds of plasmid variants that include new origins of replication (perhaps targeting groups that were less readily engineered) and antibiotic resistance or other new selection modules from parts libraries as they become available (Gilbert *et al*., 2023; Martínez-García *et al*., 2023). The use of modular Golden Gate assembly for plasmid construction makes it simple to incorporate new genetic parts into libraries (Geddes, Mendoza-Suárez and Poole, 2019; Geddes *et al*., 2024). Involving an acoustic liquid handler for preparing bacterial conjugation mixes accelerates and automates the laborious task of combining of large numbers of alternate donor strains in a single conjugation mix (Figure 1B). Similarly, expanding screening from a set of 94 microbes to hundreds or thousands of microbes is readily achievable with our workflows, designed with throughput in mind at each step. Incorporation of an acoustic liquid handler also accelerates and miniaturizes next-generation sequencing portions of the workflow (Benz *et al*., 2024).

Overall, VECTOR will facilitate future studies testing new tools for engineering diverse microbiome members and facilitate the genetic engineering of a wide variety of undomesticated strains. We expect a direct utility for labelling isolates from synthetic communities with DNA barcodes or fluorescent proteins to facilitate measurements of community assembly (Ronda *et al*., 2019; Mendoza-Suárez *et al*., 2020; Daniel *et al*., 2024; Jorrin *et al*., 2024; Ordon *et al*., 2024), and the plasmids we used herein have shown stability in the absence of antibiotic selection or are stabilized by genome integration (Geddes *et al*., 2024). The BEVA plasmids utilized in VECTOR can also readily be adapted for a wide variety of genetic engineering strategies. For example, gene editing with CRISPR (Wang *et al*., 2021), genome integration of large payloads via CRAGE (Wang *et al*., 2019), or mutagenesis with massively parallel sequencing (Liu Hualan *et al*., 2018). These approaches could be deployed using plasmid architectures identified by VECTOR^L^ screening, with the targeted engineering facilitated by VECTOR^S^ workflows.

## Acknowledgements

This work used resources of the Center for Computationally Assisted Science and Technology (CCAST) at North Dakota State University, which were made possible in part by NSF MRI Award No. 2019077. We thank the Thomas Glass Innovation Core staff (Scott Hoselton and Kaycie Schmidt) in the Department of Microbiological Sciences, North Dakota State University for technical contributions to this work.

## Data Availability

Protocols have been deposited at dx.doi.org/10.17504/protocols.io.q26g7m581gwz/v1 and http://dx.doi.org/10.17504/protocols.io.rm7vzkwr8vx1/v1

Scripts for data analysis are available at https://github.com/NDSU-Geddes-Lab/eng-potential-vector DOI: 10.5281/zenodo.15025842

Plasmids have been deposited to addgene under deposit agreement 85591

Sequence data has been deposited in the NCBI Sequence Read Archive under BioProject ID PRJNA1235777

