## Supplemental Data for "Evaluation of Engineering Potential in Undomesticated Microbes with VECTOR"

**Supplementary Information**

### Supplemental Tables

#### Supplemental Table S1. Model bacterial strains

| Strain | Description | Source |
| --- | --- | --- |
| DH5α | Competent *Escherichia coli* cells. Genotype: *F– φ80lacZΔ M15 Δ (lacZYA-argF) U169 recA1 endA1 hsdR17 (rK– mK+) phoA supE44 λ- thi–1 gyrA96 relA1* | (*DH5α Competent Cells - US*, n.d.) |
| EC100D^TM^ Pir+ | Competent *Escherichia coli* cells. Genotype: *F- mcrA Δ(mrr-hsdRMS-mcrBC) Φ80dlacZΔM15 ΔlacX74 recA1 endA1 araD139 Δ(ara, leu)7697 galU galK λ- rpsL (StrR) nupG pir+(DHFR)* | (*TransforMax^TM^ EC100D^TM^ Pir+ and Pir-116 Electrocompetent* E. Coli*, Competent Cells, Biosearch Technologies*, n.d.) |
| One Shot^TM^ Pir 1 | Competent *Escherichia coli* cells. Genotype: *F–∆lac169 rpoS(Am) robA1 creC510 hsdR514 endA recA1 uidA(∆MluI)::pir-116* | (*One Shot^TM^ PIR1 Chemically Competent* E. Coli, n.d., p. 1) |
| ST18 | Competent *Escherichia coli* cells. Genotype: *F- RP4-2(Km::Tn7,Tc::Mu-1) pro-82 LAMpir recA1 endA1 thiE1 hsdR17 creC510* *λpir ΔhemA* | (Thoma & Schobert, 2009) |
| RmP110 | *Sinorhizobium meliloti.* Wildtype derived from Rm1021 with corrected pstC allele | (Yuan et al., 2006) |
| USDA110 | *Bradyrhizobium japonicum* | (Kaneko et al., 2002) |
| R7A | *Mesorhizobium loti* | (Sullivan et al., 2013) |
| Sp7 | *Azospirillum brasilense.* Wild-type strain Sp7 (ATCC 29145). Isolated from Digitaria decumbens roots. Grass symbiont. Nitrate reducer/Nitrogen fixer. | (Magalhães et al., 1978) |
| IRBG74 | *Rhizobium spp*. Isolated from Sesbania cannabina. Forms N_2_ fixing nodules in dry and submerged conditions. Known rice endophyte, used as an innoculant to improve growth, health, yeild. | (Crook et al., 2013) |
| M5A1 | *Klebsiella oxytoca* | (Bender, 1977) |
| ORS571 | *Azorhizobium caulinodans* | (Lee et al., 2008) |

#### Supplemental Table S2. Corn and pea microbiome isolates

| ASV Name | Strain Name | ASV Name | Strain Name |
| --- | --- | --- | --- |
| P013 | *Pseudomonas thivervalensis* | C003 | *Reyranella aquatilis* |
| P017 | *Pseudomonas punonensis* | C016 | *Pseudomonas corrugata* |
| P026 | *Pseudomonas poae* | C018 | *Pseudamonas koreensis/Pseudamonas reinekei* |
| P036 | *Microbacterium arthrosphaerae* | C023 | *Acinetobacter calcoaceticus* |
| P048 | *Brevundimonas intermedia* | C029 | *Rhizobium herbae/Pararhizobium polonicum* |
| P054 | *Neorhizobium huautlense* | C031 | *Methylobacterium radiotolerans* |
| P056 | *Rhizobium skierniewicense* | C034 | *Caulobacter rhizosphaerae/Caulobacter henricii* |
| P062 | *Phyllobacterium brassicacearum* | C067 | *Pseudoxanthomonas indica/Pseudoxanthomonas mexicana* |
| P067 | *Pseudomonas peli* | C072 | *Stenotrophomonas terrae/Stenotrophomonas humi* |
| P070 | *Stutzerimonas zhaodongensis* | C074 | *Stenotrophomonas maltophilia* |
| P079 | *Dokdonella ginsengisoli* | C080 | *Lysobacter enzymogenes/Lysobacter firmicutimachus* |
| P092 | *Xanthomonas albilineans* | C085 | *Flavobacterium ginsengiterrae/Flavobacterium compostarboris* |
| P094 | *Pseudoxanthomonas wuyuanensis* | C086 | *Flavobacterium saccharophilum/Flavobacterium aquidurense* |
| P100 | *Stenotrophomonas rhizophila* | C094 | *Flavobacterium olei/Flavobacterium granuli* |
| P102 | *Bosea vestrisii* | C100 | *Pedobacter nyackensis/Pedobacter trunci* |
| P108 | *Flavobacterium hercynium* | C101 | *Pedobacter caeni/Pedobacter steynii* |
| P110 | *Flavobacterium hibisc* | C102 | *Pedobacter bambusae/Pedobacter quisquiliarum* |
| P111 | *Flavobacterium branchiicola* | C109 | *Sphingobacterium ginsenosidimutans/Sphingobacterium detergens* |
| P116 | *Flavobacterium piscis* | C117 | *Sphingobium mellinum/Sphingobium herbicidovorans* |
| P124 | *Pedobacter panaciterrae* | C122 | *Rahnella aquatilis/Rahnella inusitata* |
| P126 | *Mucilaginibacter rubeus* | C123 | *Luteibacter jiangsuensis/Luteibacter rhizovicinus* |
| P127 | *Mucilaginibacter lappiensis* | C126 | *Niabella hibiscisoli/Niabella yanshanensis* |
| P129 | *Sphingomonas mali* | C128 | *Chitinophaga rhizosphaerae/Chitinophaga caseinilytica* |
| P136 | *Sphingopyxis fribergensis* | C137 | *Chryseobacterium ginsenosidimutans/Chryseobacterium geocarposphaerae* |
| P143 | *Flavobacterium terriphilum* | C139 | *Chryseobacterium vietnamense/Chryseobacterium bernardetii* |
| P146 | *Pantoea agglomerans* | C149 | *Microbacterium thalassium/Microbacterium lacticum* |
| P150 | *Rhodanobacter glycinis* | C157 | *Paenarthrobacter nitroguajacolicus/Paenarthrobacter aurescens* |
| P154 | *Chryseobacterium soldanellicola* | C162 | *Streptomyces globisporus/Streptomyces sindenensis* |
| P158 | *Dyadobacter endophyticus* | C167 | *Cellulomonas cellasea* |
| P163 | *Pseudarthrobacter enclensis* | C169 | *Cellulosimicrobium cellulans/Cellulosimicrobium funkei* |
| P168 | *Kocuria uropygioeca* | C173 | *Paenibacillus polysaccharolyticus/Paenibacillus cucumis* |
| P169 | *Microbacterium paraoxydans* | C176 | *Mycolicibacterium tusciae* |
| P171 | *Microbacterium yannicii* | C186 | *Nocardioides kongjuensis/Nocardioides nitrophenolicus* |
| P172 | *Microbacterium liquefaciens* | C188 | *Aeromicrobium tamlense/Aeromicrobium choanae* |
| P173 | *Paenibacillus seodonensis* | C189 | *Aeromicrobium fastidiosum* |
| P177 | *Mycolicibacterium frederiksbergens* | C195 | *Variovorax paradoxus* |
| P180 | *Agromyces aureus* | C207 | *Herminiimonas saxobsidens/Herminiimonas contaminans* |
| P189 | *Pseudomonas batumici* | C210 | *Achromobacter xylosoxidans/Achromobacter denitrificans* |
| P194 | *Variovorax Paradoxus* | C212 | *Duganella zoogloeoides* |
| P198 | *Limnohabitans parvus* | C218 | *Galbitalea soli* |
| P204 | *Paraburkholderia graminis* | C223 | *Rathayibacter caricis/Rathayibacter tritici* |
| P205 | *Achromobacter mucicolens* | C224 | *Conyzicola lurida/Conyzicola nivalis* |
| P218 | *Pseudorhodoferax soli* | C228 | *Curtobacterium oceanosedimentum* |
| P221 | *Hydrogenophaga borbori* | C230 | *Pseudoclavibacter terrae/Pseudoclavibacter helvolus* |
| P233 | *Methylophilus flavus* | C232 | *Polaromonas aquatica/Polaromonas jejuensis* |
| P236 | *Rhodococcus fascians* | C248 | *Psychrobacillus psychrodurans/Psychrobacillus psychrotolerans* |
| P241 | *Priestia megaterium* | C250 | *Bacillus australimaris/Bacillus safensis* |

#### Supplemental Table S3. Primers for genetic engineering

| Primer Name | Sequence (5’-3’) | Purpose | Length (bp) | Tm (°C) | % GC |
| --- | --- | --- | --- | --- | --- |
| RWND005 | AAAGGTCTCTGGAGCGTCTCAACTACCTAGGCGGCCTTAATTAA | Mariner Cassette 1 Forward | 44 | 54.3 | 47.7 |
| RWND006 | TTTGGTCTCAACCTTCTCCGAAGCGTGGAAA | Mariner Cassette 1 Reverse | 32 | 55.6 | 48.4 |
| RWND007 | TTTGGTCTCAAGGTTCACCGGTCGCT | Mariner Cassette 2 Forward | 26 | 55.6 | 53.8 |
| RWND008 | TTTGGTCTCAATGGCGTCTCATCTGGATTTTGAGACACAAGACGTC | Mariner Cassette 2 Reverse | 46 | 55.2 | 45.7 |
| RWND014 | TTGGAAAGAGTCTTTTGGCT | pBBR1 Forward | 20 | 53 | 40 |
| RWND016 | ATTCAAGCCAAGAACAAGC | pBBR1 Reverse | 19 | 52 | 42 |
| RWND017 | TTCCACATCGAGGAAGAAAA | RSF1010 Forward | 20 | 52 | 40 |
| RWND019 | CGGATGTTATCGACCAGTA | RSF1010 Reverse | 19 | 52 | 47.4 |
| RWND020 | ACAGGTTGGATGATAAGTCC | pMarC9-R6K Forward | 20 | 53 | 45 |
| RWND022 | GCCCTAGATCTGGATTTTGA | pMarC9-R6K Reverse | 20 | 53 | 45 |
| RWND026 | TTCCTCGTCGATCAGGACCT | RK2 Forward | 20 | 59 | 55 |
| RWND027 | CGAGCTGCGGGCCGACGATG | RK2 Reverse | 20 | 69 | 75 |

#### Supplemental Table S4. Plasmids for Golden Gate cloning

| Vector Name | Backbone | Purpose | Module Size (bp) | 5’-side | 3’-side | Source |
| --- | --- | --- | --- | --- | --- | --- |
| B0032m_BC | DVA | MoClo Basic part. Ribosome binding site, modified from Bba_0032. | 2108 | TACT | AATG | (Iverson et al., 2016) |
| J23106_AB | DVA | MoClo Basic part. Constitutive promoter. Anderson Series. | 2108 | GGAG | TACT | (Iverson et al., 2016) |
| pNDGG001 | pLVC-P2-neo | Neomycin/Kanamycin resistance module. | 1044 | GCAA | ACTA | (Geddes et al., 2024) |
| pNDGG003 | L1-RK2-Tc-par | GG assembly from pOGG010, pOGG012, pOGG014, pOGG042, and pOGG004 | 7081 | CGCT | GGAG | (Geddes et al., 2024) |
| pNDGG004 | L1-pBBR1-Tc-par | GG assembly from pOGG011, pOGG012, pOGG014, pOGG042, and pOGG004 | 6400 | CGCT | GGAG | (Geddes et al., 2024) |
| pNDGG005 | pGGASelect | RSF1010 origin of replication, *oriT* from RK2*.* | 2159 | ACTA | TTAC | (Geddes et al., 2024) |
| pNDGG0049 | L1-pBBR1-Gm-par | GG assembly from pOGG004, pOGG009, pOGG011, pOGG012, and pOGG014 | 5229 | CGCT | GGAG | (Geddes et al., 2024) |
| pNDGG0050 | L1-RK2-Gm-par | GG assembly from pOGG004, pOGG009, pOGG010, pOGG012, and pOGG014 | 5930 | CGCT | GGAG | (Geddes et al., 2024) |
| pNDGG0053 | L1-pBBR1-Km-par | GG assembly from pOGG004, pNDGG001, pOGG011, pOGG012, and pOGG014 | 5460 | CGCT | GGAG | (Geddes et al., 2024) |
| pNDGG054 | L1-RK2-Km-par | GG assembly from pOGG004, pNDGG001, pOGG010, pOGG012, and pOGG014 | 6161 | CGCT | GGAG | (Geddes et al., 2024) |
| pNDGG056 | L1-RSF1010-Gm-par | GG assembly from pGQ0015, pOGG009, pNDGG005, pOGG012, and pOGG014 | 9765 | CGCT | GGAG | (Geddes et al., 2024) |
| pNDGG057 | L1-RSF1010-Tc-par | GG assembly from pOGG004, pOGG042, pNDGG005, pOGG012, and pOGG014 | 10916 | CGCT | GGAG | (Geddes et al., 2024) |
| pNDGG059 | L1-RSF1010-Km-par | GG assembly from pOGG004, pNDGG001, pNDGG005, pOGG012, and pOGG014 | 9996 | CGCT | GGAG | (Geddes et al., 2024) |
| pNDGG066 | L1- pMarc9-R6k-Gm | GG assembly from pOGG004, pOGG009, pNDGG071, and pOGG014 | 4254 | CGCT | GGAG | This study |
| pNDGG067 | L1- pMarc9-R6k-Km | GG assembly from pOGG004, pNDGG001, pNDGG071, and pOGG014 | 4486 | CGCT | GGAG | This study |
| pNDGG068 | L1- pMarc9-R6k-Tc | GG assembly from pOGG004, pOGG042, pNDGG071, and pOGG014 | 5406 | CGCT | GGAG | This study |
| pNDGG071 | pGGASelect | Amplified pMarC9-R6K regions, flanked with BsmBI recognition sites | 3214 | ACTA | TTAC | This study |
| pOGG004 | pLVC-P1 | Golden Gate Level 1 cloning site. Contains *lacZ*. | 723 | TGCC | GCAA | (Geddes et al., 2019) |
| pOGG009 | pLVC-P2-gent | Gentamicin resistance module. | 813 | GCAA | ACTA | (Geddes et al., 2019) |
| pOGG010 | pLVC-P3-RK2 | RK2 origin of replication, *oriT* from RK2*.* | 2485 | ACTA | TTAC | (Geddes et al., 2019) |
| pOGG011 | pLVC-P3-pBBR1 | pBBR1 origin of replication, *oriT* from RK2*.* | 1784 | ACTA | TTAC | (Geddes et al., 2019) |
| pOGG012 | pLVC-P4-par | Par stability accessory module. | 2408 | TTAC | CAGA | (Geddes et al., 2019) |
| pOGG013 | pLVC-ELT3 | End linker for position 4, circularizes linear assembly. | 113 | TTAC | TGCC | (Geddes et al., 2019) |
| pOGG014 | pLVC-ELT4 | End linker for position 5, circularizes linear assembly. | 113 | CAGA | TGCC | (Geddes et al., 2019) |
| pOGG037 | pL0M-SC-sfGFP | Super-folder GFP green fluorescence reporter. | 722 | AATG | GCTT | (Geddes et al., 2019) |
| pOGG042 | pLVC-P2-Tet | Tetracycline resistance module. | 1964 | GCAA | ACTA | (Geddes et al., 2019) |

#### Supplemental Table S5. Plasmids generated by Golden Gate cloning

| Vector Name | Description | Vector size (bp) | Oligo Name | Concentration [pmol/µl] | % GC | Sequence (5'->3') |
| --- | --- | --- | --- | --- | --- | --- |
| pNDMS450 | GG assembly from pNDGG003, J23106_AB, B0032m_BC, pOGG037, and Barcode 104. | 7899 | 806rcbc104_sense | 100 | 51 | GCTTATCTACCGAAGCCGT  TTTACAACGTCGTGACTGGG |
|  |  |  | 806rcbc104_anti | 100 | 53 | AGCGCCCAGTCACGACGTT  GTAAAACGGCTTCGGTAGAT |
| pNDMS451 | GG assembly from pNDGG004, J23106_AB, B0032m_BC, pOGG037, and Barcode 402. | 7198 | 806rcbc402_sense | 100 | 53 | GCTTACCGGAGTAGGACGT  TTTACAACGTCGTGACTGGG |
|  |  |  | 806rcbc402_anti | 100 | 56 | AGCGCCCAGTCACGACGTT  GTAAAACGTCCTACTCCGGT |
| pNDMS452 | GG assembly from pNDGG049, J23106_AB, B0032m_BC, pOGG037, and Barcode 403. | 6947 | 806rcbc403_sense | 100 | 51 | GCTTTGAGGACTACCTCGT  TTTACAACGTCGTGACTGGG |
|  |  |  | 806rcbc403_anti | 100 | 53 | AGCGCCCAGTCACGACGTT  GTAAAACGAGGTAGTCCTCA |
| pNDMS453 | GG assembly from pNDGG050, J23106_AB, B0032m_BC, pOGG037, and Barcode 404. | 6748 | 806rcbc404_sense | 100 | 53 | GCTTCAATCGGCTTGCCGT  TTTACAACGTCGTGACTGGG |
|  |  |  | 806rcbc404_anti | 100 | 56 | AGCGCCCAGTCACGACGTT  GTAAAACGGCAAGCCGATTG |
| pNDMS456 | GG assembly from pNDGG053, J23106_AB, B0032m_BC, pOGG037, and Barcode 407. | 6278 | 806rcbc407_sense | 100 | 51 | GCTTGGAGGAGCAATACGT  TTTACAACGTCGTGACTGGG |
|  |  |  | 806rcbc407_anti | 100 | 53 | AGCGCCCAGTCACGACGTT  GTAAAACGTATTGCTCCTCC |
| pNDMS457 | GG assembly from pNDGG054, J23106_AB, B0032m_BC, pOGG037, and Barcode 408. | 6979 | 806rcbc408_sense | 100 | 53 | GCTTAGCGACGAAGACCGT  TTTACAACGTCGTGACTGGG |
|  |  |  | 806rcbc408_anti | 100 | 56 | AGCGCCCAGTCACGACGTT  GTAAAACGGTCTTCGTCGCT |
| pNDMS458 | GG assembly from pNDGG056, J23106_AB, B0032m_BC, pOGG037, and Barcode 410. | 10583 | 806rcbc410_sense | 100 | 53 | GCTTTGGAAGAACGGCCGT  TTTACAACGTCGTGACTGGG |
|  |  |  | 806rcbc410_anti | 100 | 56 | AGCGCCCAGTCACGACGTT  GTAAAACGGCCGTTCTTCCA |
| pNDMS459 | GG assembly from pNDGG057, J23106_AB, B0032m_BC, pOGG037, and Barcode 413. | 10814 | 806rcbc413_sense | 100 | 48 | GCTTGATATACCAGTGCGT  TTTACAACGTCGTGACTGGG |
|  |  |  | 806rcbc413_anti | 100 | 51 | AGCGCCCAGTCACGACGTT  GTAAAACGCACTGGTATATC |
| pNDMS461 | GG assembly from pNDGG059, J23106_AB, B0032m_BC, pOGG037, and Barcode 418. | 11734 | 806rcbc418_sense | 100 | 51 | GCTTTAGTGTCGGATCCGT  TTTACAACGTCGTGACTGGG |
|  |  |  | 806rcbc418_anti | 100 | 53 | AGCGCCCAGTCACGACGTT  GTAAAACGGATCCGACACTA |
| pNDMS462 | GG assembly from pNDGG066, J23106_AB, B0032m_BC, pOGG037, and Barcode 414. | 6224 | 806rcbc414_sense | 100 | 48 | GCTTAACAAACTGCCACGT  TTTACAACGTCGTGACTGGG |
|  |  |  | 806rcbc414_anti | 100 | 51 | AGCGCCCAGTCACGACGTT  GTAAAACGTGGCAGTTTGTT |
| pNDMS463 | GG assembly from pNDGG067, J23106_AB, B0032m_BC, pOGG037, and Barcode 415. | 5304 | 806rcbc415_sense | 100 | 48 | GCTTGTAGACATGTGTCGT  TTTACAACGTCGTGACTGGG |
|  |  |  | 806rcbc415_anti | 100 | 51 | AGCGCCCAGTCACGACGTT  GTAAAACGACACATGTCTAC |
| pNDMS464 | GG assembly from pNDGG068, J23106_AB, B0032m_BC, pOGG037, and Barcode 416. | 6054 | 806rcbc416_sense | 100 | 51 | GCTTTACAGTTACGCGCGT  TTTACAACGTCGTGACTGGG |
|  |  |  | 806rcbc416_anti | 100 | 53 | AGCGCCCAGTCACGACGTT  GTAAAACGCGCGTAACTGTA |

#### Supplemental Table S6. Bioinformatics scripts

| File Category | Program | File/Script Name |
| --- | --- | --- |
| Echo-Protocols | Echo Cherry Pick | CornandPeaTrial_Cherrypick.xlsx |
| Echo-Protocols | Echo Cherry Pick | CornandPeaTrial_SinglePlasmid_Cherrypick.xlsx |
| Echo-Protocols | Echo Cherry Pick | MockConjugation_Cherrypick.xlsx |
| Echo-Protocols | Echo Cherry Pick | MockConjugation_MultiPlasmid_Cherrypick.xlsx |
| Growth_Curve_Scripts | R-Studio | Growth_Curve_Analysis.R |
| Growth_Curve_Scripts | R-Studio | Growth_Curve_Analysis_Engineered.R |
| Growth_Curve_Scripts | R-Studio | MockConjugation_OD_GrowthCurve.Rmd |
| Growth_Curve_Scripts | R-Studio | MockConjugation_RFU_GrowthCurve.Rmd |
| Growth_Curve_Scripts | R-Studio | Multiplate_Control_Cytation5_Cleanup.Rmd |
| Growth_Curve_Scripts | R-Studio | Multiplate_Cytation5_Cleanup.Rmd |
| Plasmid_ID_Scripts | Jupyter | Plasmid_ID.ipynb |
| Plasmid_ID_Scripts | R-Studio | Plasmid_ID_Cleanup.Rmd |

#### Supplemental Table S7. Primary PCR primers for well barcoding

| Forward Primer Number | Index Sequence (5’-3’) | Reverse Primer Name | Index Sequence (5’-3’) |
| --- | --- | --- | --- |
| 1_F | TCGTCGGCAGCGTCAGATGTGTATAAG  AGACAGGCTACATCACGCATGGTATGGA | A_R | GTCTCGTGGGCTCGGAGATGTGTATAAGAG  ACAGGCTCCCCAGTCACGACGTTGTAAAACG |
| 2_F | TCGTCGGCAGCGTCAGATGTGTATAAGA  GACAGTGTGTCATCACGCATGGTATGGA | B_R | GTCTCGTGGGCTCGGAGATGTGTATAAGAGA  CAGCTAGTCCCAGTCACGACGTTGTAAAACG |
| 3_F | TCGTCGGCAGCGTCAGATGTGTATAAGA  GACAGAGTCTGCATCACGCATGGTATGGA | C_R | GTCTCGTGGGCTCGGAGATGTGTATAAGAGA  CAGTAGATCCCCAGTCACGACGTTGTAAAACG |
| 4_F | TCGTCGGCAGCGTCAGATGTGTATAAG  AGACAGATCACATCACGCATGGTATGGA | D_R | GTCTCGTGGGCTCGGAGATGTGTATAAGAG  ACAGTCGCCCCAGTCACGACGTTGTAAAACG |
| 5_F | TCGTCGGCAGCGTCAGATGTGTATAAGA  GACAGGACGACATCACGCATGGTATGGA | E_R | GTCTCGTGGGCTCGGAGATGTGTATAAGAGA  CAGCCTTACCCAGTCACGACGTTGTAAAACG |
| 6_F | TCGTCGGCAGCGTCAGATGTGTATAAGA  GACAGTCGTCGCATCACGCATGGTATGGA | F_R | GTCTCGTGGGCTCGGAGATGTGTATAAGAGA  CAGCATAACCCCAGTCACGACGTTGTAAAACG |
| 7_F | TCGTCGGCAGCGTCAGATGTGTATAAG  AGACAGTGCTCATCACGCATGGTATGGA | G_R | GTCTCGTGGGCTCGGAGATGTGTATAAGAGA  CAGCAGACCCAGTCACGACGTTGTAAAACG |
| 8_F | TCGTCGGCAGCGTCAGATGTGTATAAGA  GACAGCAGTTCATCACGCATGGTATGGA | H_R | GTCTCGTGGGCTCGGAGATGTGTATAAGAGA  CAGTGTTCCCCAGTCACGACGTTGTAAAACG |
| 9_F | TCGTCGGCAGCGTCAGATGTGTATAAGA  GACAGACATGTCATCACGCATGGTATGGA |  |  |
| 10_F | TCGTCGGCAGCGTCAGATGTGTATAAGA  GACAGGCGGCATCACGCATGGTATGGA |  |  |
| 11_F | TCGTCGGCAGCGTCAGATGTGTATAAGA  GACAGGTTGACATCACGCATGGTATGGA |  |  |
| 12_F | TCGTCGGCAGCGTCAGATGTGTATAAGA  GACAGGTGGCTCATCACGCATGGTATGGA |  |  |

#### Supplemental Table S8. Secondary PCR primers for plate indexing

| Reverse Primer Name | Index Sequence (5’-3’) | Forward Primer Name | Index Sequence (5’-3’) |
| --- | --- | --- | --- |
| D501 | AATGATACGGCGACCACCGAGATCTA  CACTATAGCCTTCGTCGGCAGCGTC | A701 | CAAGCAGAAGACGGCATACGAGA  TGTCGTGATGTCTCGTGGGCTCGG |
| D502 | AATGATACGGCGACCACCGAGATCTA  CACATAGAGGCTCGTCGGCAGCGTC | D701 | CAAGCAGAAGACGGCATACGAGA  TCGAGTAATGTCTCGTGGGCTCGG |
| D503 | AATGATACGGCGACCACCGAGATCTA  CACCCTATCCTTCGTCGGCAGCGTC | D702 | CAAGCAGAAGACGGCATACGAGA  TTCTCCGGAGTCTCGTGGGCTCGG |
| D504 | AATGATACGGCGACCACCGAGATCTA  CACGGCTCTGATCGTCGGCAGCGTC | D703 | CAAGCAGAAGACGGCATACGAGA  TAATGAGCGGTCTCGTGGGCTCGG |
| D505 | AATGATACGGCGACCACCGAGATCTA  CACAGGCGAAGTCGTCGGCAGCGTC | D704 | CAAGCAGAAGACGGCATACGAGA  TGGAATCTCGTCTCGTGGGCTCGG |
| D506 | AATGATACGGCGACCACCGAGATCTA  CACTAATCTTATCGTCGGCAGCGTC | D705 | CAAGCAGAAGACGGCATACGAGA  TTTCTGAATGTCTCGTGGGCTCGG |
| D507 | AATGATACGGCGACCACCGAGATCTA  CACCAGGACGTTCGTCGGCAGCGTC | D706 | CAAGCAGAAGACGGCATACGAGA  TACGAATTCGTCTCGTGGGCTCGG |
| D508 | AATGATACGGCGACCACCGAGATCTA  CACGTACTGACTCGTCGGCAGCGTC | D707 | CAAGCAGAAGACGGCATACGAGA  TAGCTTCAGGTCTCGTGGGCTCGG |
| A501 | AATGATACGGCGACCACCGAGATCTA  CACTGAACCTTTCGTCGGCAGCGTC | D708 | CAAGCAGAAGACGGCATACGAGA  TGCGCATTAGTCTCGTGGGCTCGG |
| N501 | AATGATACGGCGACCACCGAGATCTA  CACTAGATCGCTCGTCGGCAGCGTC | D709 | CAAGCAGAAGACGGCATACGAGA  TCATAGCCGGTCTCGTGGGCTCGG |
| N502 | AATGATACGGCGACCACCGAGATCTA  CACCTCTCTATTCGTCGGCAGCGTC | D710 | CAAGCAGAAGACGGCATACGAGA  TTTCGCGGAGTCTCGTGGGCTCGG |
| N503 | AATGATACGGCGACCACCGAGATCTA  CACTATCCTCTTCGTCGGCAGCGTC | D711 | CAAGCAGAAGACGGCATACGAGA  TGCGCGAGAGTCTCGTGGGCTCGG |
| N504 | AATGATACGGCGACCACCGAGATCTA  CACAGAGTAGATCGTCGGCAGCGTC | N701 | CAAGCAGAAGACGGCATACGAGA  TTCGCCTTAGTCTCGTGGGCTCGG |
| N505 | AATGATACGGCGACCACCGAGATCTA  CACGTAAGGAGTCGTCGGCAGCGTC | N702 | CAAGCAGAAGACGGCATACGAGA  TCTAGTACGGTCTCGTGGGCTCGG |
| N506 | AATGATACGGCGACCACCGAGATCTA  CACACTGCATATCGTCGGCAGCGTC | N703 | CAAGCAGAAGACGGCATACGAGA  TTTCTGCCTGTCTCGTGGGCTCGG |
| N507 | AATGATACGGCGACCACCGAGATCTA  CACAAGGAGTATCGTCGGCAGCGTC | N704 | CAAGCAGAAGACGGCATACGAGA  TGCTCAGGAGTCTCGTGGGCTCGG |
|  |  | N705 | CAAGCAGAAGACGGCATACGAGA  TAGGAGTCCGTCTCGTGGGCTCGG |
|  |  | N706 | CAAGCAGAAGACGGCATACGAGA  TCATGCCTAGTCTCGTGGGCTCGG |
|  |  | N707 | CAAGCAGAAGACGGCATACGAGA  TGTAGAGAGGTCTCGTGGGCTCGG |
|  |  | N708 | CAAGCAGAAGACGGCATACGAGA  TCCTCTCTGGTCTCGTGGGCTCGG |
|  |  | D712 | CAAGCAGAAGACGGCATACGAGA  TCTATCGCTGTCTCGTGGGCTCGG |
|  |  | N710 | CAAGCAGAAGACGGCATACGAGA  TCAGCCTCGGTCTCGTGGGCTCGG |
|  |  | N711 | CAAGCAGAAGACGGCATACGAGA  TTGCCTCTTGTCTCGTGGGCTCGG |
|  |  | N712 | CAAGCAGAAGACGGCATACGAGA  TTCCTCTACGTCTCGTGGGCTCGG |

#### Supplemental Table S9. Differences in growth (OD_600_) between experimental and control wells in test diazotrophs.

| Strain | Mean Diff. | Adjusted P Value | Mean Diff. | Adjusted P Value | Mean Diff. | Adjusted P Value |
| --- | --- | --- | --- | --- | --- | --- |
| **Gentamicin** | **20μg/ml** | | **50μg/ml** | | **100μg/ml** | |
| RmP110 | 0.3643 | 0.0277 | 0.5063 | 0.0457 | 0.1253 | 0.2409 |
| R7A | 0.297 | 0.0885 | 0.5007 | 0.0487 | 0.1117 | 0.3501 |
| Sp7 | 0.1813 | 0.5037 | 0.4987 | 0.0499 | 0.1663 | 0.0681 |
| IRBG74 | 0.3243 | 0.0554 | 0.6319 | 0.0109 | 0.2333 | 0.0078 |
| M5A1 | 0.4273 | 0.0094 | 0.604 | 0.0149 | 0.3883 | <0.0001 |
| ORS751 | 0.403 | 0.0142 | 0.6982 | 0.0052 | 0.2483 | 0.0049 |
| **Neomycin** | **20μg/ml** | | **50μg/ml** | | **100μg/ml** | |
| RmP110 | 0.3742 | 0.0452 | 0.4444 | 0.0004 | 0.4234 | 0.0018 |
| R7A | 0.327 | 0.0936 | 0.422 | 0.0007 | 0.3877 | 0.0038 |
| Sp7 | 0.2932 | 0.1551 | 0.4019 | 0.001 | 0.3576 | 0.007 |
| IRBG74 | 0.3911 | 0.0347 | 0.4086 | 0.0009 | 0.5777 | 0.0001 |
| M5A1 | 0.4636 | 0.0113 | 0.4957 | 0.0002 | 0.443 | 0.0013 |
| ORS751 | 0.4178 | 0.0229 | 0.4188 | 0.0007 | 0.518 | 0.0003 |
| **Tetracycline** | **5μg/ml** | | **10μg/ml** | | **20μg/ml** | |
| RmP110 | 0.4303 | 0.0009 | 0.2637 | 0.0002 | 0.4669 | 0.0106 |
| R7A | 0.3947 | 0.0018 | 0.2157 | 0.0012 | 0.4307 | 0.0186 |
| Sp7 | 0.225 | 0.0835 | 0.133 | 0.0403 | 0.3967 | 0.0316 |
| IRBG74 | 0.01367 | >0.9999 | 0.07067 | 0.4993 | 0.5423 | 0.0034 |
| M5A1 | 0.008333 | >0.9999 | 0.158 | 0.0132 | 0.5896 | 0.0017 |
| ORS751 | 0.08633 | 0.8766 | 0.04533 | 0.8701 | 0.5371 | 0.0037 |

### Supplemental Figures

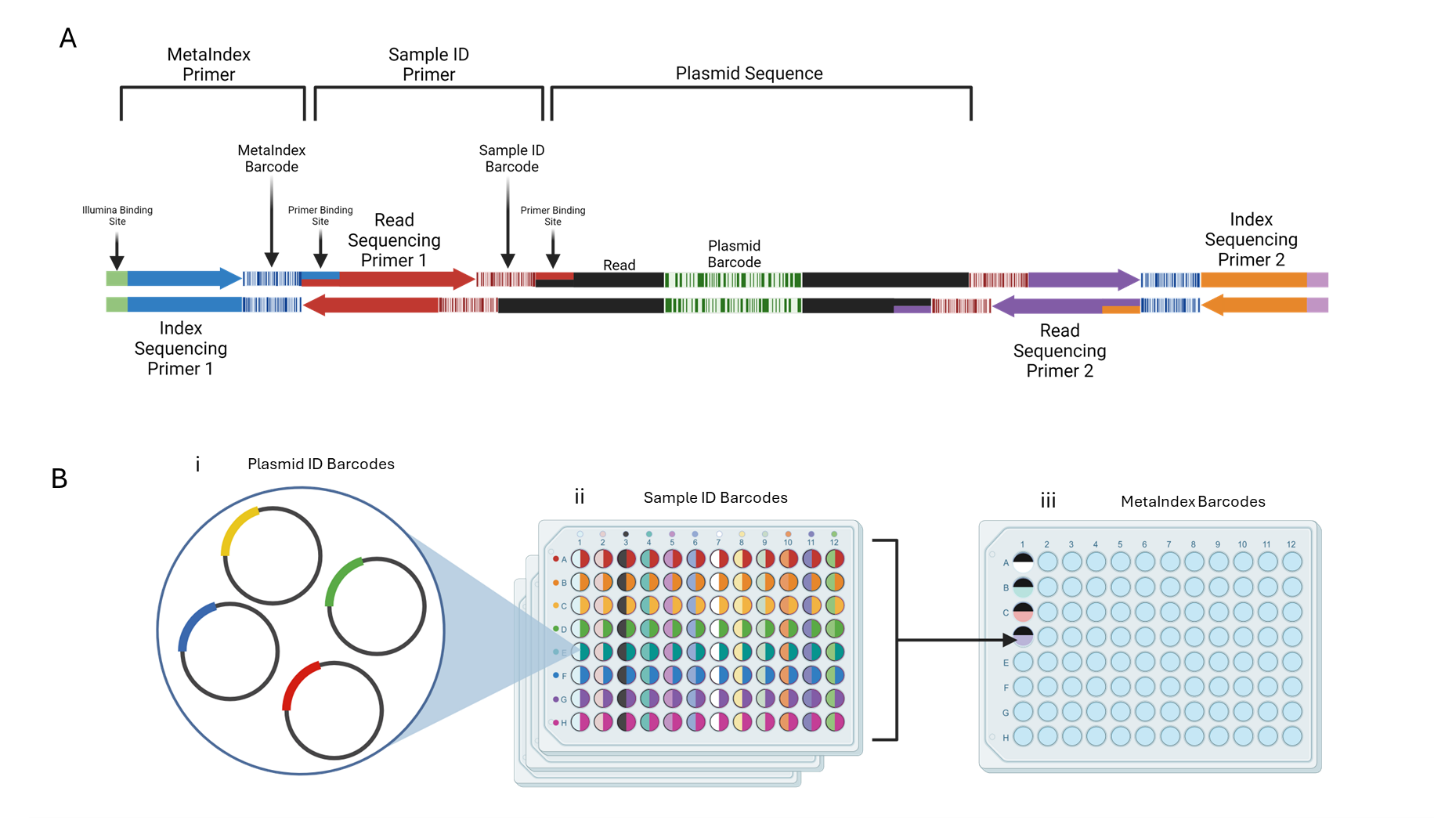

Supplemental Figure 1. Multi-layered dual indexing strategy for next-generation sequencing. **(A)** Schematic representation of the structure of plasmid sequences after the second round of PCR (PCR2). **(B)** Barcode sequencing workflow for identifying plasmids. (i) DNA from lysed samples is distributed into individual wells. (ii) Each well is labeled through a primary PCR using a dual well-ID barcode added with specific primers. The color-coded primer combinations illustrate how 12 forward primers are distributed across columns, while 8 reverse primers are assigned to rows. (iii) The contents of each plate are pooled into a single well of a new plate and labeled with a dual meta-indexing PCR. This second PCR incorporates Nextera barcodes and adaptors for Illumina sequencing. Color examples represent different primer and barcode combinations.

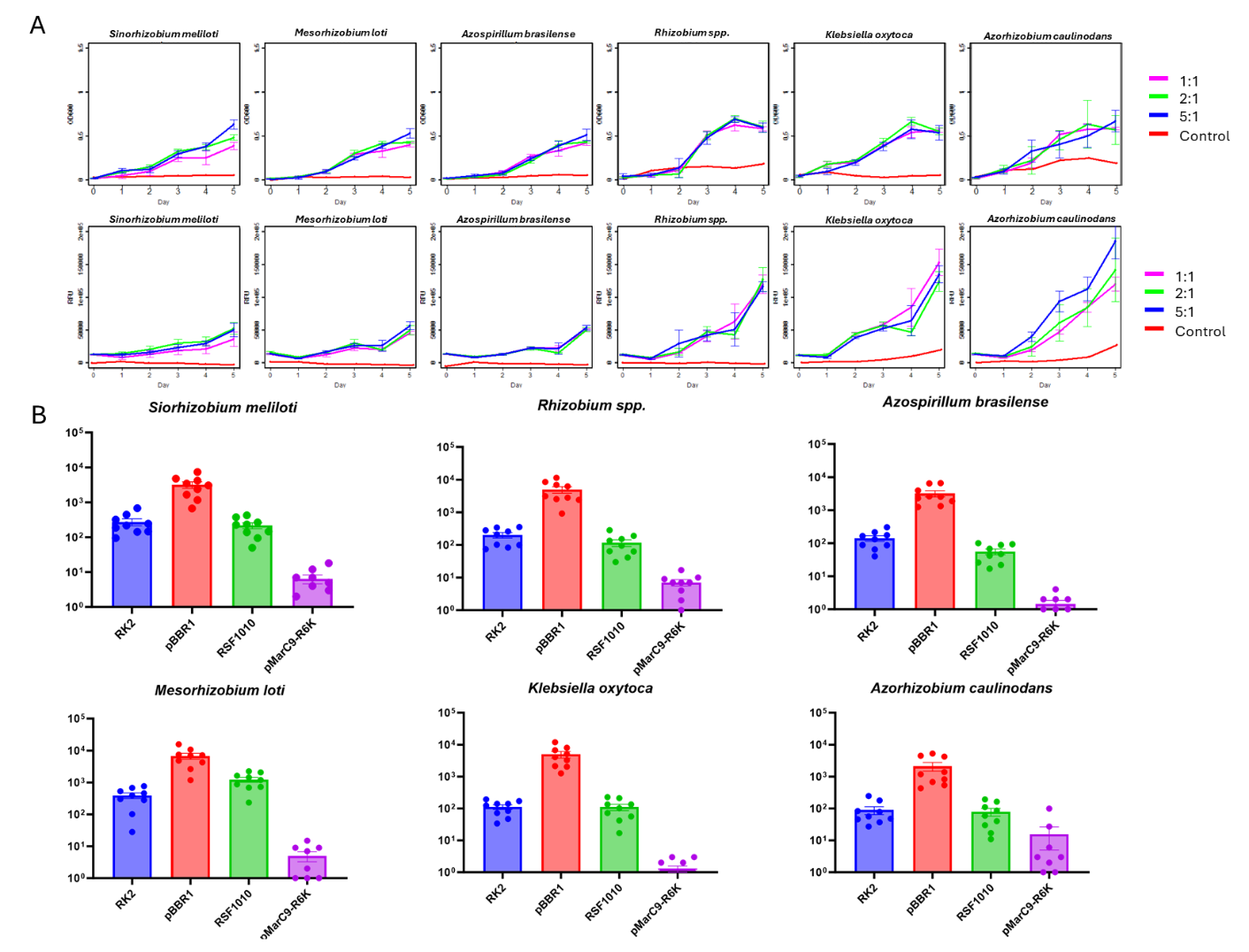

Supplemental Figure S2. Growth curves and plasmid sequencing reads from mock community. **(A)** Mock community growth curves showing the progression of OD600 (above) and RFU (below) over five days, grown in Tetracycline (20μg/ml) supplemented TY broth. Each growth curve displays data from trials with varying recipient to donor ratios, indicated in the keys to right. Lines were constructed using the mean value of each replicate measure for each day. **(B)** Plots indicating the number of sequencing reads of each plasmid in our library among the members of the mock community of microbes. Each plot point represents the number of reads identified as the corresponding *ori* below*.*

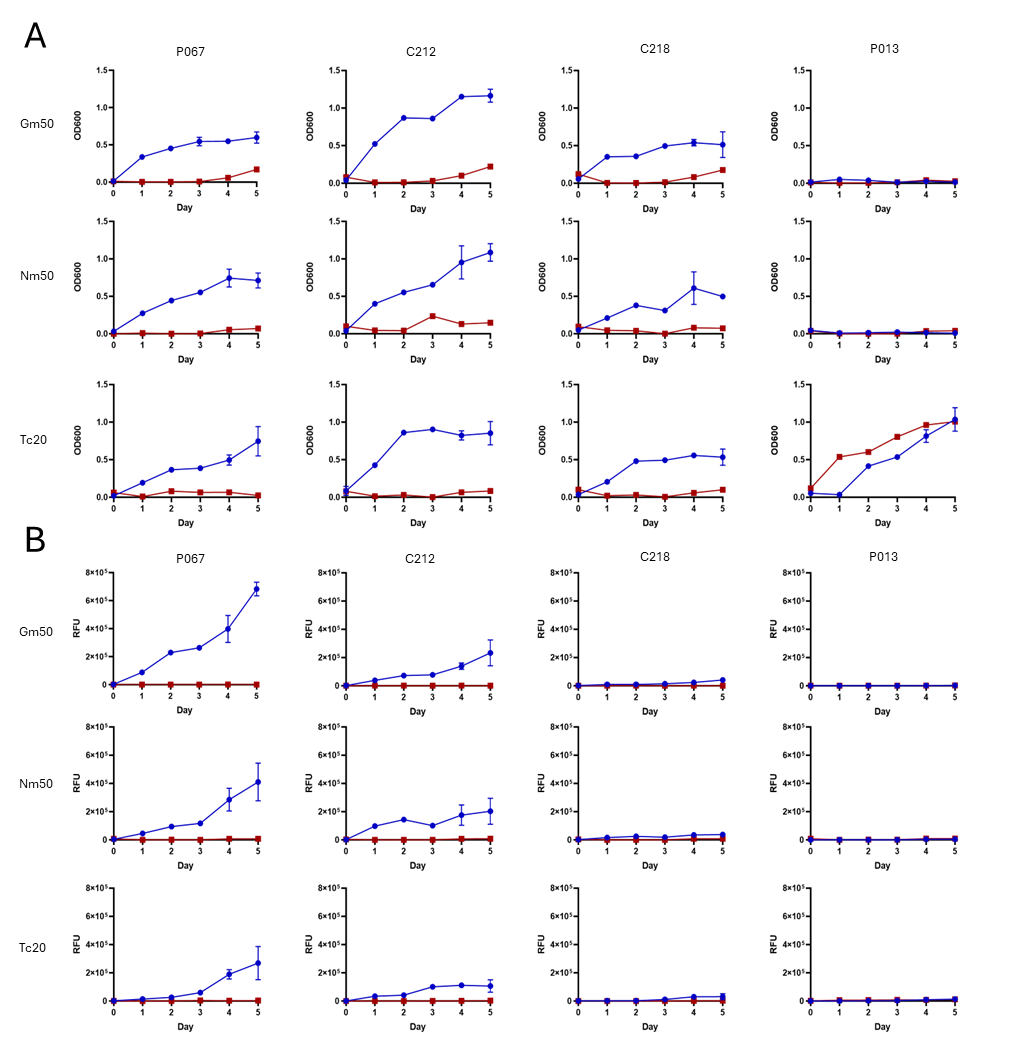

Supplemental Figure S3. Example growth and fluorescent curves from corn/pea microbiome members. Five-day liquid growth curves of the four strains shown as representatives of engineered strain growth (Figure 2A). The ASV number of each strain is listed at the top of each column and antibiotic condition is listed to the left of each row. Engineered samples are shown in blue with plot points representing the mean value of triplicate measures, with standard error bars, and the control samples shown in red. Growth curves show both **(A)** OD600 and **(B)** RFU, plotted on a log10 scale.

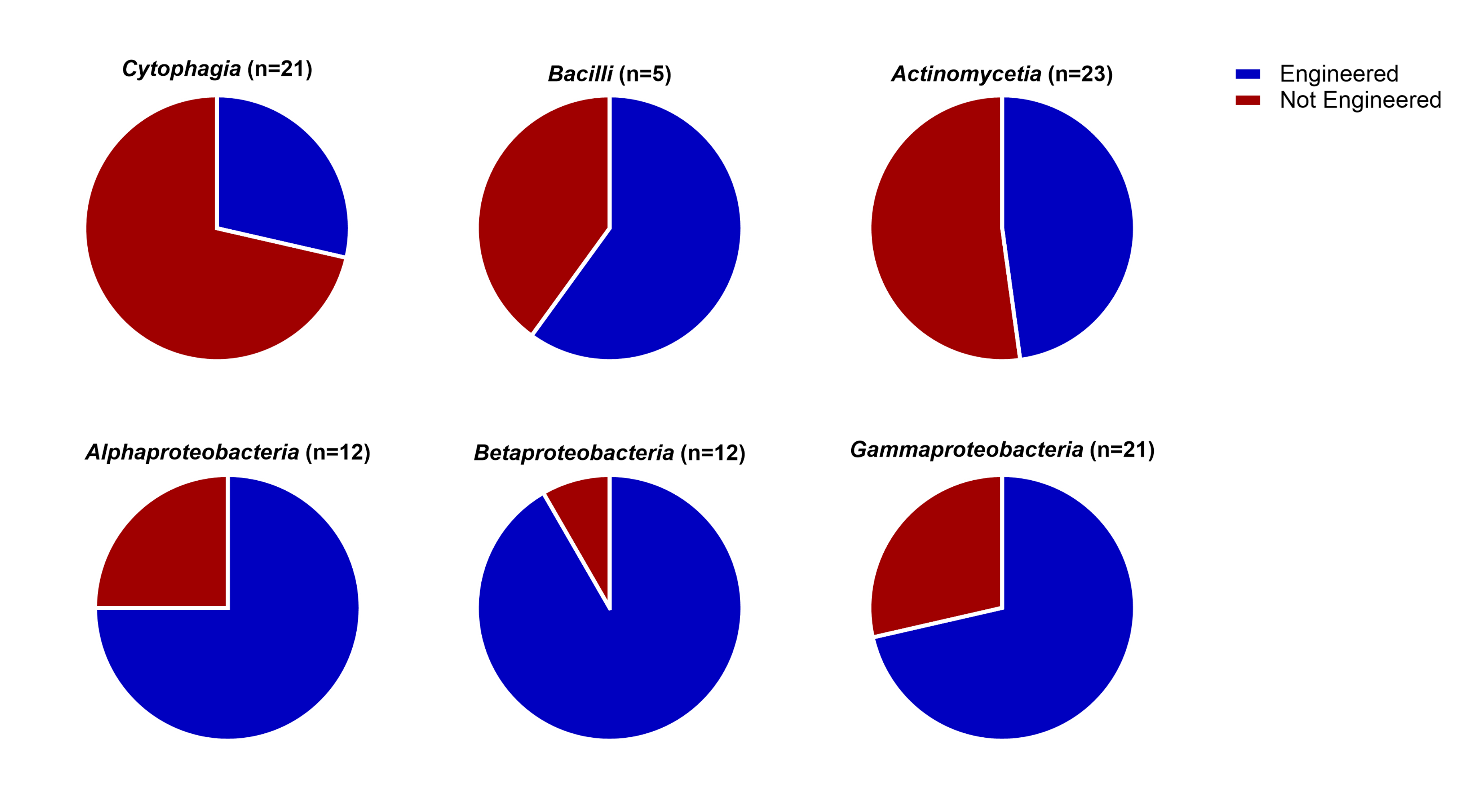

Supplemental Figure S4. Pie charts showing the proportion of engineered versus non-engineered strains across various phylogenetic classes. Each chart is labeled with the corresponding class name and the number of strains within that class. Engineering designation is based on results from streak plating on selection media, where a strain is classified as engineered if it forms fluorescent colonies under at least one of three antibiotic conditions and maintains the morphology of its parental strain.

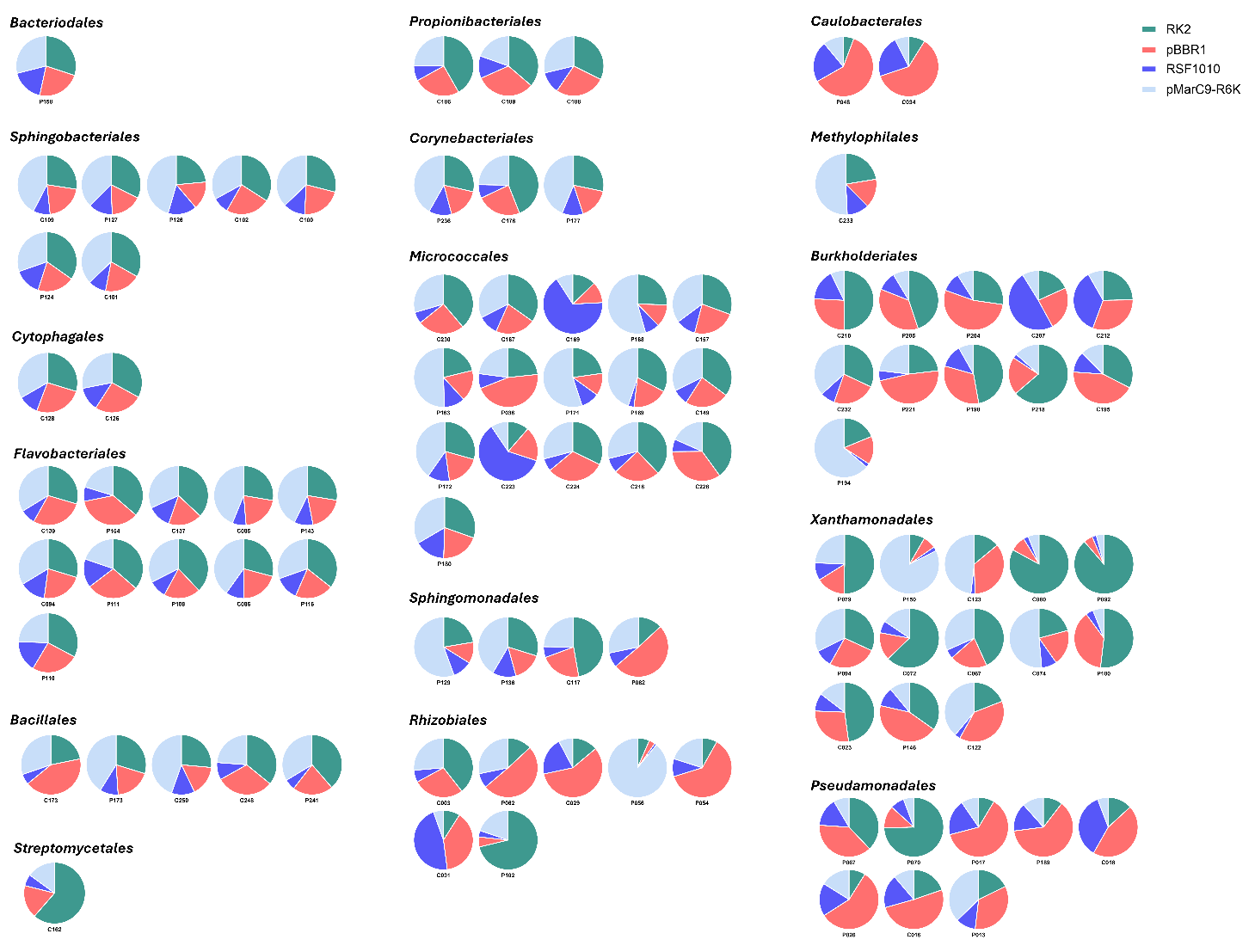

Supplemental Figure 5. Pie charts illustrating the normalized relative read frequencies of each origin of replication (ori) within individual strains. The percentages were adjusted to account for plasmid copy number. Strains are labeled with their ASV numbers below each chart and grouped by phylogenetic order, indicated above each group. The pie charts represent the distribution of the four *oris*: RK2 (green), pBBR1 (pink), RSF1010 (blue), and pMarC9-R6K (light blue).

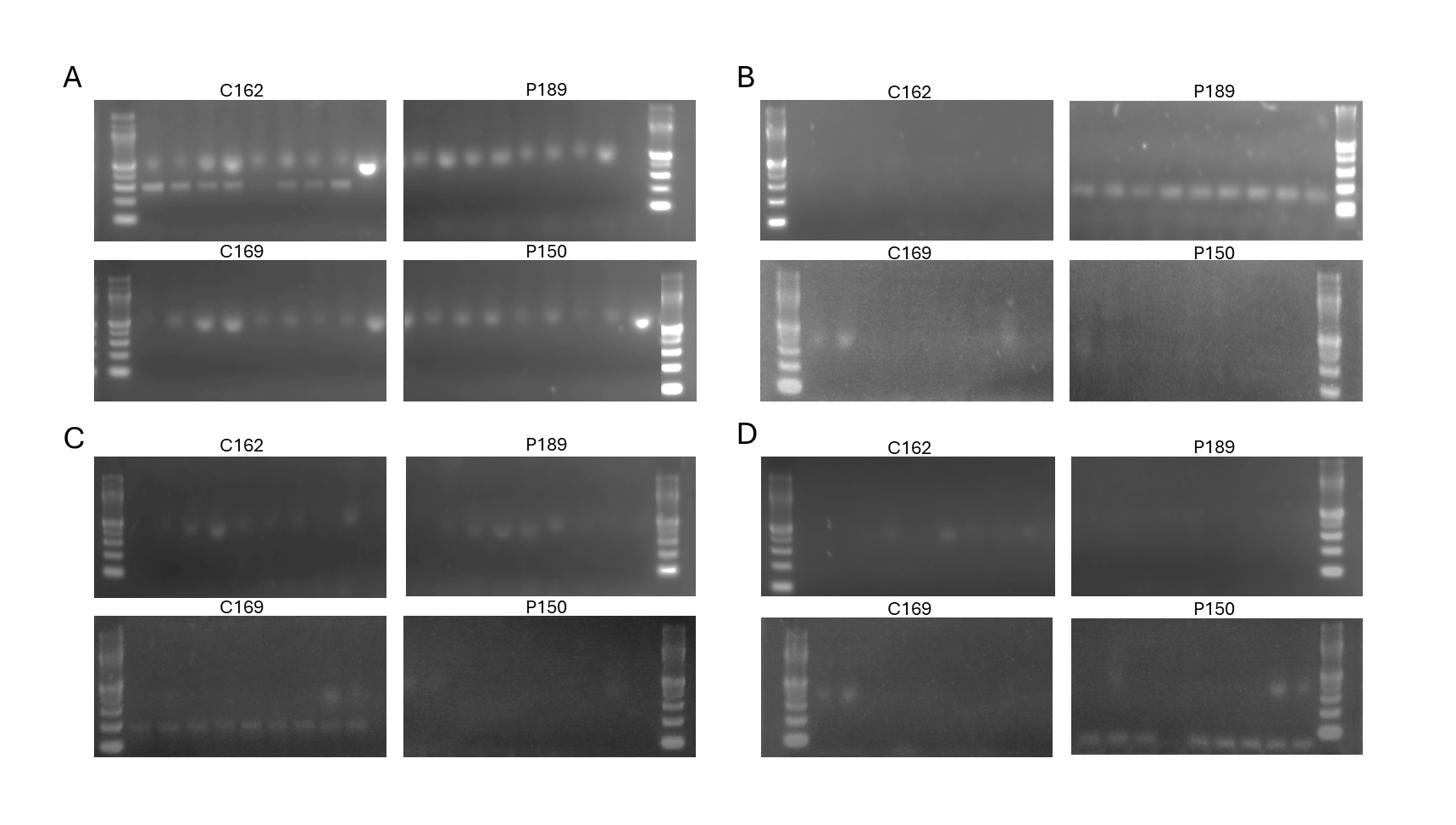

Supplemental Figure S6. Electrophoresis gels displaying PCR results using plasmid-specific primers to amplify DNA from strains C162, P189, C169, and P150. Each primer set was designed to target a specific region of the corresponding plasmid, producing the expected band sizes: RK2 (*trfA*) – 300 bp, pBBR1 (*oriV*) – 170 bp, RSF1010 (*oriV*) – 188 bp, and pMarC9-R6K (between Mosaic inverted repeat and terminator 1) – 61 bp. Gel images are organized by the plasmid origin being screened in each strain: **(A)** RK2, **(B)** pBBR1, **(C)** RSF1010, and **(D)** pMarC9-R6K.

### Supplementary Methods

#### Data Analysis for Figure Preparation.

To generate Figure 1D, daily measurements from the Cytation 5 were reorganized to simplify analysis and merge replicate values. The raw output files were processed using the Multiplate_Cytation5_Cleanup.Rmd script for experimental plates and the Multiplate_Control_Cytation5_Cleanup.Rmd script for control plates. The processed data, consisting of OD_data and RFU_data, were compiled into a new Excel file and analyzed using the Growth_Curve_Analysis.R script. This workflow was repeated for each antibiotic condition to ensure consistency.

Figures 2C-E were created using the same Excel file generated during the processing of Cytation 5 data. The script Growth_Curve_Analysis_Engineered.R was run to analyze the reorganized data. An excel file was used to select specific data relevant to the engineered strain experiments, enabling detailed comparisons under the specified conditions.

To generate Figures 2F and 2G, plasmid reads were adjusted to account for expected sequencing coverage per plate. Each plate's reads were scaled to a total of 100,000, eliminating sequencing biases across replicates. For analysis, only reads corresponding to each plasmid's specific antibiotic selection condition (e.g., RK2 plasmids with acc3 analyzed from gentamicin plates) were averaged to determine plasmid presence in each strain. To compare plasmid reads across different origins of replication (*oris*), the data were normalized by plasmid copy number. The read counts were divided by the following copy numbers: 7 for RK2 (Kües and Stahl, 1989), 30 for pBBR1 (Jiang *et al.*, 2017), 12 for RSF1010 (Meyer, 2009), and 1 for pMarC9-R6K transposon. The final normalized data were categorized by ori and antibiotic resistance module and visualized as heatmaps using GraphPad Prism. This data was also used to create Supplemental Figures S4 and S5.

To generate Supplemental Table S9, the growth of each strain on the final day was compared to its corresponding negative control. An ANOVA test followed by Šídák’s multiple comparisons test was performed to calculate the mean difference and adjusted p-value for each strain at each antibiotic concentration.

To generate the growth curves in Supplemental Figure S2, daily measurements from the Cytation 5 were reorganized, merged, and plotted using two distinct scripts. For OD600 growth curves, the script MockConjugation_OD_GrowthCurve.Rmd was used to reshape the daily measurement files. The initial portion of the script generated an intermediate file, which was further processed manually into a final file named Mock_OD_Growth.csv. The remainder of the script utilized this file to create a .png file containing six growth curve plots for the specified antibiotic condition. This process was repeated for all antibiotic conditions, and the resulting plots were overlaid to produce six comprehensive growth curve plots, including all conditions and controls. A similar workflow was followed to create the RFU plots. The script MockConjugation_RFU_GrowthCurve.Rmd was used to reshape the RFU data into a file named Mock_RFU_Growth.csv. This file was then used to generate the final RFU growth curve plots in the same format.

Supplemental Figure S3 was created by extracting the mean OD and RFU values, along with the standard error for each strain, on each antibiotic and for each day. These values were obtained from the daily outputs of the Multiplate_Cytation5_Cleanup.Rmd and Multiplate_Control_Cytation5_Cleanup.Rmd scripts. The extracted mean and standard error values were used to plot the growth curves in GraphPad Prism.

### Supplementary Datasets

#### Supplementary Dataset A. Logistic regression models based on ∆OD and ∆RFU

Logistic regression model summaries, *p*-values for χ^2^ goodness-of-fit tests, and residual plots are included below for each antibiotic: Gentamicin (Gm50), Neomycin (Nm50), and Tetracycline (Tc20). With ∆OD and ∆RFU as predictors, all three models were considered good fits to the data, with both predictors yielding significant effects. A table of these results was included in the main manuscript. Verbatim outputs from R and diagnostic residual plots are included here for reference.

**Gm50 (∆OD and ∆RFU)**

Call:

glm(formula = gm50_eng ~ delta_od + delta_rfu, family = "binomial",

data = filter(wide_data_gm50, time == 6))

Coefficients:

Estimate Std. Error z value Pr(>|z|)

(Intercept) -3.3090	0.8749 -3.782 0.000155 ***

delta_od	4.6084	1.1977	3.848 0.000119 ***

delta_rfu	2.8642	0.8117	3.529 0.000418 ***

---

Signif. codes: 0 ‘***’ 0.001 ‘**’ 0.01 ‘*’ 0.05 ‘.’ 0.1 ‘ ’ 1

(Dispersion parameter for binomial family taken to be 1)

Null deviance: 129.928 on 93 degrees of freedom

Residual deviance: 73.621 on 91 degrees of freedom

AIC: 79.621

Number of Fisher Scoring iterations: 6

> pchisq(mod_gm50$deviance, mod_gm50$df.residual, lower.tail = F)

[1] 0.9083258

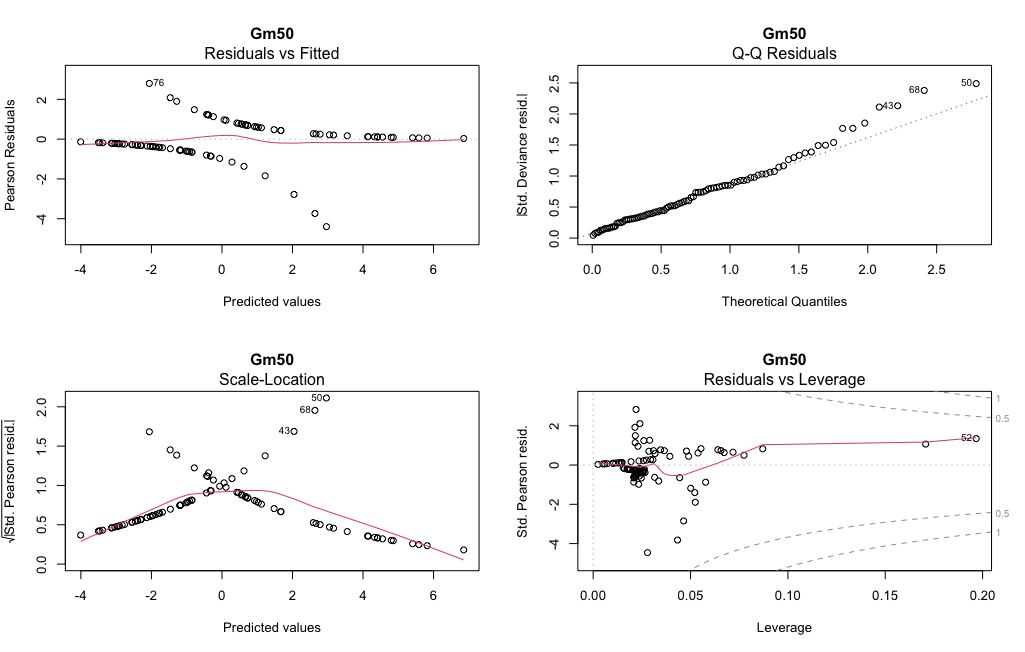

**Nm50 (∆OD and ∆RFU)**

Call:

glm(formula = nm50_eng ~ delta_od + delta_rfu, family = "binomial",

data = filter(wide_data_nm50, time == 6))

Coefficients:

Estimate Std. Error z value Pr(>|z|)

(Intercept) -1.9988	0.5921 -3.376 0.000736 ***

delta_od	3.0850	0.8837	3.491 0.000482 ***

delta_rfu	1.7981	0.6990	2.572 0.010100 *

---

Signif. codes: 0 ‘***’ 0.001 ‘**’ 0.01 ‘*’ 0.05 ‘.’ 0.1 ‘ ’ 1

(Dispersion parameter for binomial family taken to be 1)

Null deviance: 129.630 on 93 degrees of freedom

Residual deviance: 90.031 on 91 degrees of freedom

AIC: 96.031

Number of Fisher Scoring iterations: 5

> pchisq(mod_nm50$deviance, mod_nm50$df.residual, lower.tail = F)

[1] 0.5090111

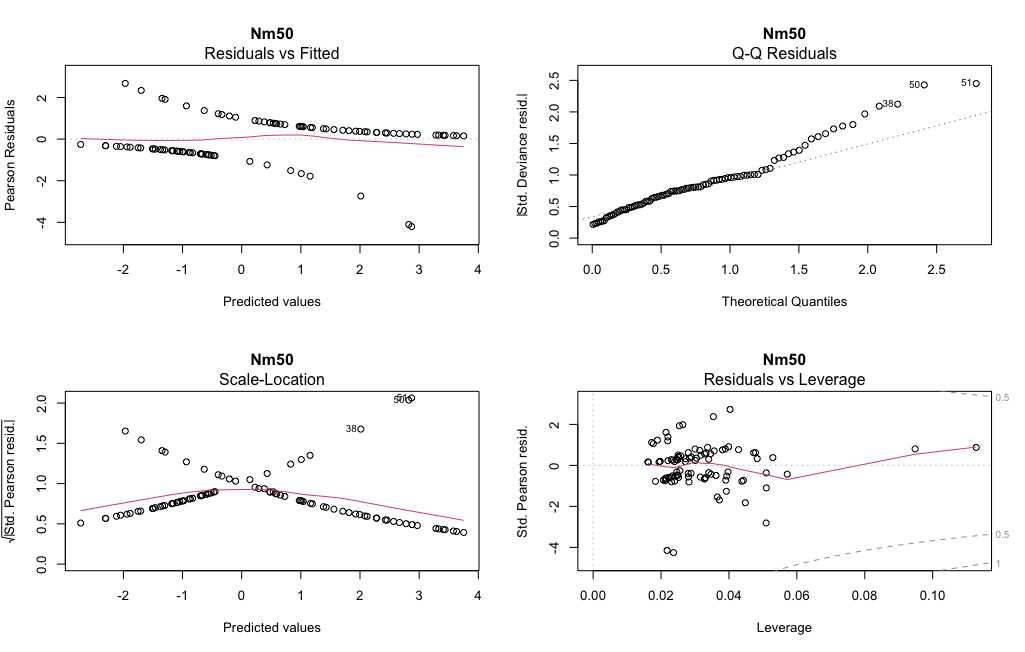

**Tc20 (∆OD and ∆RFU)**

Call:

glm(formula = tc20_eng ~ delta_od + delta_rfu, family = "binomial",

data = filter(wide_data_tc20, time == 6))

Coefficients:

Estimate Std. Error z value Pr(>|z|)

(Intercept) -2.9753	0.6578 -4.523 6.09e-06 ***

delta_od	1.7872	0.8342	2.142	0.0322 *

delta_rfu	3.2077	0.7791	4.117 3.84e-05 ***

---

Signif. codes: 0 ‘***’ 0.001 ‘**’ 0.01 ‘*’ 0.05 ‘.’ 0.1 ‘ ’ 1

(Dispersion parameter for binomial family taken to be 1)

Null deviance: 128.219 on 93 degrees of freedom

Residual deviance: 94.148 on 91 degrees of freedom

AIC: 100.15

Number of Fisher Scoring iterations: 4

> pchisq(mod_tc20$deviance, mod_tc20$df.residual, lower.tail = F)

[1] 0.3897285

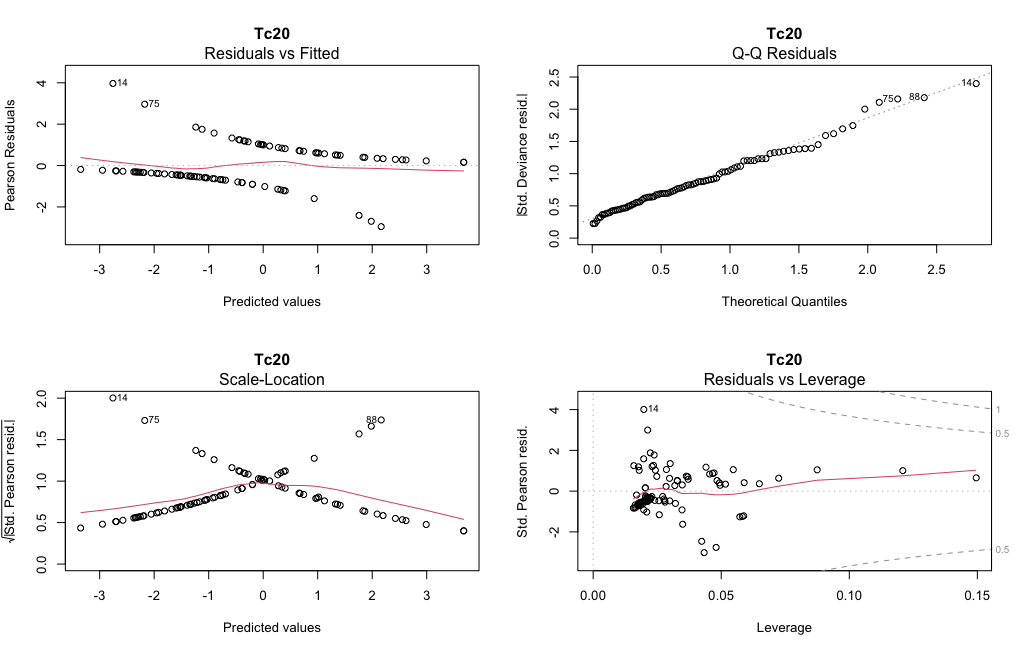

#### Supplementary Dataset B: Logistic regression models based on taxonomic class

Logistic regression model summaries, *p*-values for χ^2^ goodness-of-fit tests, and residual plots are included below for each antibiotic: Gentamicin (Gm50), Neomycin (Nm50), and Tetracycline (Tc20). With taxonomic class labels as predictors, model fits were much poorer (χ^2^ *p*-values < 0.10), and most class effects were non-significant. Therefore, taxonomy does not appear to have as much predictive power for engineerability as do the observed ∆OD and ∆RFU for each strain. For these reasons, results tables for the taxonomic class models were not included in the main manuscript, but have been included here for reference.

**Gm50 (taxonomic class)**

Call:

glm(formula = gm50_eng ~ 0 + class, family = "binomial", data = filter(wide_data_gm50,

time == 6))

Coefficients:

Estimate Std. Error z value Pr(>|z|)

classActinomycetia	-0.4418	0.4272 -1.034 0.30107

classAlphaproteobacteria	1.0986	0.6667	1.648 0.09937 .

classBacilli	0.4055	0.9129	0.444 0.65692

classBetaproteobacteria	1.0986	0.6667	1.648 0.09937 .

classCytophagia	-2.2513	0.7433 -3.029 0.00246 **

classGammaproteobacteria	0.2877	0.4410	0.652 0.51414

---

Signif. codes: 0 ‘***’ 0.001 ‘**’ 0.01 ‘*’ 0.05 ‘.’ 0.1 ‘ ’ 1

(Dispersion parameter for binomial family taken to be 1)

Null deviance: 130.31 on 94 degrees of freedom

Residual deviance: 106.40 on 88 degrees of freedom

AIC: 118.4

Number of Fisher Scoring iterations: 4

> pchisq(mod_gm50_class$deviance, mod_gm50_class$df.residual, lower.tail = F)

[1] 0.08847708

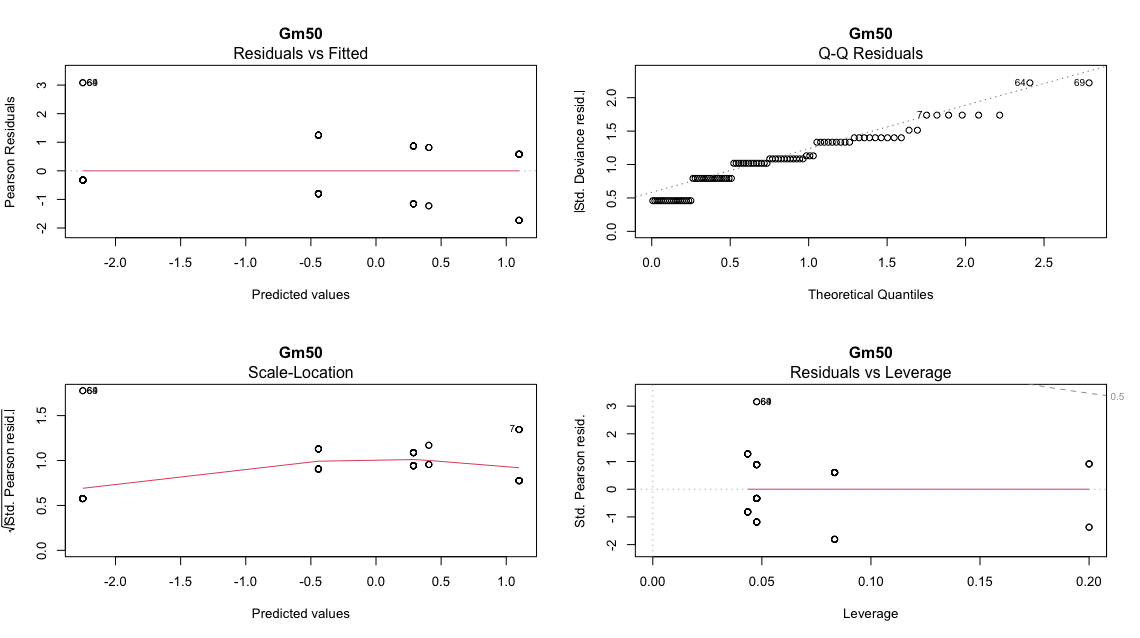

**Nm50 (taxonomic class)**

Call:

glm(formula = nm50_eng ~ 0 + class, family = "binomial", data = filter(wide_data_nm50,

time == 6))

Coefficients:

Estimate Std. Error z value Pr(>|z|)

classActinomycetia	-0.08701	0.41742 -0.208	0.8349

classAlphaproteobacteria 1.09861	0.66667	1.648	0.0994 .

classBacilli	0.40547	0.91287	0.444	0.6569

classBetaproteobacteria	2.39789	1.04403	2.297	0.0216 *

classCytophagia	-1.16315	0.51235 -2.270	0.0232 *

classGammaproteobacteria 0.28768	0.44096	0.652	0.5141

---

Signif. codes: 0 ‘***’ 0.001 ‘**’ 0.01 ‘*’ 0.05 ‘.’ 0.1 ‘ ’ 1

(Dispersion parameter for binomial family taken to be 1)

Null deviance: 130.31 on 94 degrees of freedom

Residual deviance: 110.69 on 88 degrees of freedom

AIC: 122.69

Number of Fisher Scoring iterations: 4

> pchisq(mod_nm50_class$deviance, mod_nm50_class$df.residual, lower.tail = F)

[1] 0.05142826

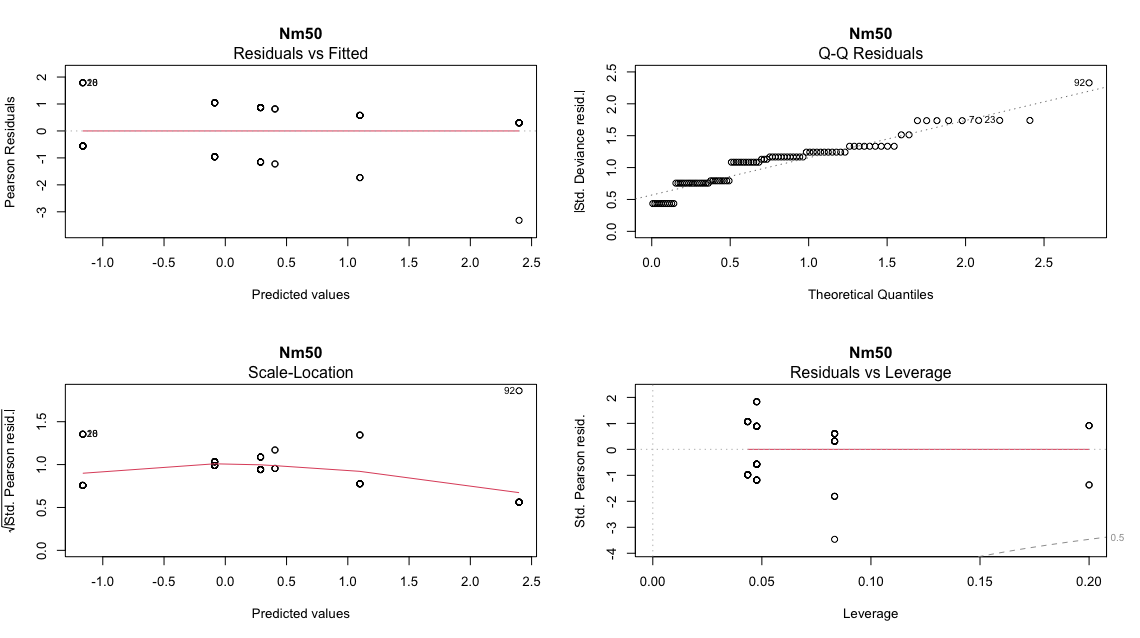

**Tc20 (taxonomic class)**

Call:

glm(formula = tc20_eng ~ 0 + class, family = "binomial", data = filter(wide_data_tc20,

time == 6))

Coefficients:

Estimate Std. Error z value Pr(>|z|)

classActinomycetia	-0.44183	0.42725 -1.034 0.30107

classAlphaproteobacteria 1.09861	0.66667	1.648 0.09937 .

classBacilli	-1.38629	1.11803 -1.240 0.21500

classBetaproteobacteria	0.69315	0.61237	1.132 0.25767

classCytophagia	-2.25129	0.74329 -3.029 0.00246 **

classGammaproteobacteria 0.09531	0.43693	0.218 0.82732

---

Signif. codes: 0 ‘***’ 0.001 ‘**’ 0.01 ‘*’ 0.05 ‘.’ 0.1 ‘ ’ 1

(Dispersion parameter for binomial family taken to be 1)

Null deviance: 130.31 on 94 degrees of freedom

Residual deviance: 106.84 on 88 degrees of freedom

AIC: 118.84

Number of Fisher Scoring iterations: 4

> pchisq(mod_tc20_class$deviance, mod_tc20_class$df.residual, lower.tail = F)

[1] 0.08392246

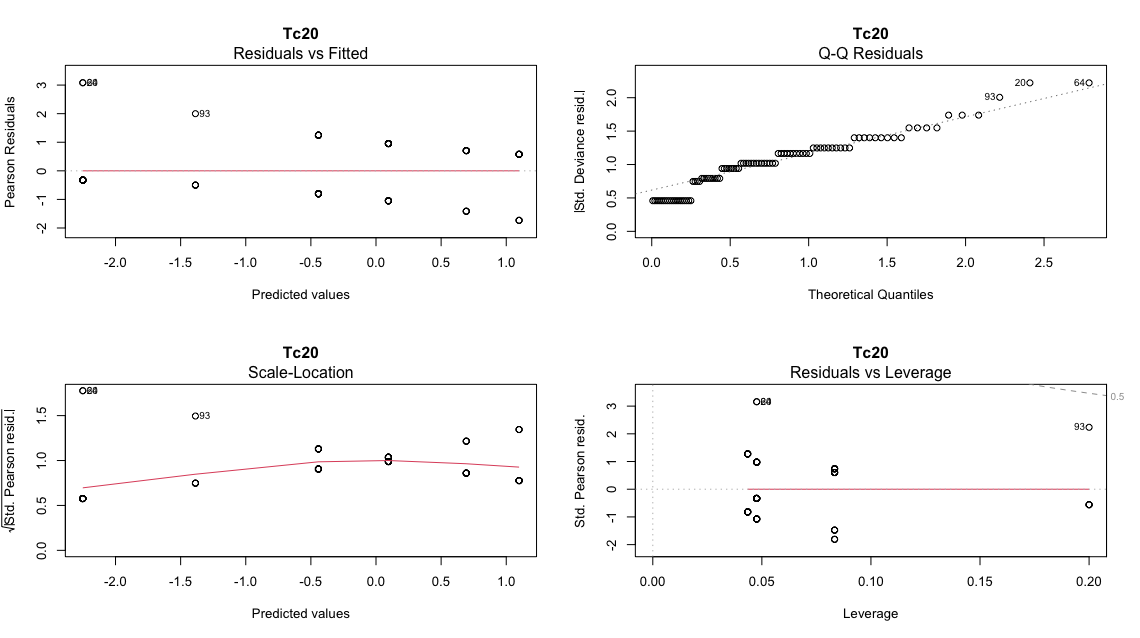
